# Pregnancy Impacts Maternal Sleep: Focus on Kynurenine Pathway Activation in Female Rats

**DOI:** 10.64898/2026.09.14.751544

**Authors:** Courtney J. Wright, Maria V. Piroli, J. Hunter Cox, Homayoun Valafar, Norma Frizzell, Ana Pocivavsek

**Affiliations:** Department of Pharmacology, Physiology, and Neuroscience, University of South Carolina Floyd School of Medicine, Columbia, SC 29209, USA; Carolina Autism and Neurodevelopment Research Center, University of South Carolina, Columbia, SC 29201, USA; Department of Computer Science and Engineering, University of South Carolina Molinaroli College of Engineering and Computing, Columbia, SC 29201, USA; Artificial Intelligence Institute, University of South Carolina, Columbia SC 29201, USA

**Keywords:** Maternal sleep, kynurenic acid, REM sleep, perinatal health, tryptophan metabolism, saliva biomarker

## Abstract

**Study Objectives:** Poor maternal sleep is associated with adverse health outcomes for women and their offspring, and perturbations to tryptophan metabolism through the kynurenine pathway (KP) may contribute. Kynurenic acid (KYNA), a neuroactive KP metabolite and endogenous modulator of glutamatergic and cholinergic neurotransmission, is of particular interest because elevated KYNA has been linked to altered sleep-wake behavior and neurodevelopmental outcomes. Because sleep patterns and KP metabolism change during pregnancy, we tested whether pregnancy modifies the effects of kynurenine supplementation on sleep-wake architecture.

**Methods:** To model enhanced KP metabolism, we used the embryonic kynurenine (EKyn) model, in which the maternal diet is supplemented with kynurenine, the initial KP metabolite, during the last week of gestation. Using a within-subject design, we evaluated sleep-wake parameters in pregnant and non-pregnant female rats and assessed KYNA in brain and saliva.

**Results:** Pregnancy increased dark phase NREM and REM sleep in control-fed (ECon) dams, a pattern largely maintained in EKyn dams. However, near parturition, EKyn dams exhibited reduced light phase REM sleep that was not observed in controls. In non-pregnant females, kynurenine supplementation produced a stronger phenotype, consistently reducing light phase REM sleep across treatment days. During pregnancy, kynurenine-associated sleep effects were detectable only when compared to within-subject non-pregnant baseline. Salivary KYNA distinguished kynurenine-fed females from matched controls and correlated with brain KYNA levels.

**Conclusion:** These findings show that pregnancy reshapes sleep-wake architecture and attenuates, but does not eliminate, kynurenine-associated REM sleep disruption, implicating KP metabolism and potentially elevated brain KYNA in maternal sleep regulation.

**Statement of Significance:** Sleep disruption during pregnancy is common and clinically important, but metabolic pathways that shape maternal sleep remain poorly defined. Using a rodent model, this study advances the concept that pregnancy changes how the brain responds to metabolic challenges, including altered tryptophan metabolism via the kynurenine pathway, which has been implicated in pregnancy complications and neurodevelopmental risk. The findings support maternal sleep as a dynamic physiologic phenotype rather than a static outcome and suggest that individualized sleep patterns may reveal disturbances missed by group comparisons alone. By linking a clinically relevant metabolic pathway to maternal sleep regulation in rats, this work provides a translational framework for future studies identifying biomarkers and strategies to protect maternal and offspring health.

## Introduction

Sleep is a biologically conserved process essential to daily functioning and overall health. During pregnancy, sleep becomes particularly critical in supporting maternal physiology and fetal development. However, this period is marked by notable sleep disturbances [1,2]. Poor maternal sleep is associated with adverse health outcomes for both women and their offspring [3-10]. Clinical studies therefore highlight the importance of restorative sleep during pregnancy; nonetheless, further research is necessary to differentiate normal sleep variations from pathophysiological disruptions during the perinatal period and to elucidate the biological mechanisms that induce or predict maternal sleep disturbances [11,12]. Rodent models are essential preclinical tools for this goal. Like humans, rats and mice experience changes in sleep patterns during the perinatal period [13-16], and experimental disruption to maternal sleep in rodents consistently produces behavioral and biochemical endophenotypes in offspring that align with adverse outcomes observed in human studies of maternal sleep disruption [17-20]. Additionally, rodent models have provided insight into significant physiological changes in maternal neurobiology that occur during peripartum and following maternal sleep disturbance [21-26]. These altered neurophysiological processes may represent clinically relevant mechanisms contributing to poor maternal and child health outcomes.

Tryptophan metabolism through the kynurenine pathway (KP) is significantly elevated during pregnancy, and this elevation serves important roles for neurodevelopment [27-32]. Further perturbations to gestational tryptophan metabolites in maternal plasma and the placenta are clinically associated with preeclampsia, fetal growth restriction, miscarriage, and maternal perinatal depressive symptoms, highlighting the importance of balanced KP metabolism during gestation for maternal and child health [32-40]. Among the neuroactive KP metabolites, kynurenic acid (KYNA) is of particular interest for neurodevelopment and maternal sleep, as it endogenously inhibits glutamatergic and cholinergic signaling [41,42], neurotransmitter systems critical to these physiological processes [43,44]. During gestation, elevated KP metabolism and enhanced KYNA levels can result from infection, inflammation, stress, and sleep disturbance can increase KP, which are risk factors for impaired offspring neurodevelopment [22,32,45-47]. Rodent offspring exposed to elevated KP metabolism during neurodevelopment exhibit long-lasting adverse developmental outcomes, including persistent behavioral alterations that pharmacological inhibition of kynurenine aminotransferase II (KAT II) can ameliorate in adulthood, supporting a mechanistic role for KYNA in these effects [48-61]. Because elevated KYNA also disrupts sleep quality and architecture [62,63], abnormal prenatal KYNA elevations may contribute to both maternal sleep disturbances and altered neurodevelopmental outcomes in offspring.

Our previous studies have demonstrated that the embryonic kynurenine (EKyn) paradigm, which models KP activation by supplementing the maternal diet with kynurenine during the last week of gestation, produces lasting biochemical and behavioral deficits in offspring that align with clinically relevant endophenotypes of neurodevelopmental disorders [49-52,54,56-59]. While the homeostatic sleep drive increases during normal pregnancy, the consequences of prenatal metabolic insults for maternal sleep architecture and sleep stability remain poorly understood. Given that both sleep patterns and KP metabolism change during normal pregnancy, the present study tested whether pregnancy modifies the effects of increased brain KYNA on sleep. Specifically, we predicted that elevated brain KYNA resulting from a kynurenine-supplemented diet would alter sleep-wake architecture and that pregnancy would influence the magnitude and pattern of KYNA-associated sleep disruptions.

To test this hypothesis, we maintained both non-pregnant and pregnant female rats on control or kynurenine-supplemented diets. We assessed biochemical and sleep-related parameters at baseline, during kynurenine exposure, and after a five-day washout period. During pregnancy, these periods corresponded to a non-pregnant baseline, the last week of pregnancy, and the postpartum period, respectively. We monitored sleep-wake behavior with implantable telemetry devices that recorded electroencephalogram (EEG), electromyography (EMG), core body temperature, and relative cage activity. We assessed sleep-wake behavior using the architecture of wakefulness, non-rapid eye movement (NREM) sleep, and rapid eye movement (REM) sleep, as well as NREM delta power, REM theta power, sleep onset, and transitions among these vigilance states. Finally, we assessed whether salivary KYNA could distinguish between treatment conditions and serve as a peripheral correlate of elevated brain KYNA. We found that pregnancy significantly alters female NREM and REM sleep patterns and delays the reduction in light phase REM sleep from kynurenine diet, though kynurenine effects were detectable only when compared to within-subject non-pregnant baseline sleep. Salivary KYNA levels distinguished kynurenine-fed females from matched controls and demonstrated a significant positive correlation with brain KYNA levels. Together, these findings identify a unique role for brain KYNA in sleep-wake homeostasis during gestation and highlight the importance of individual sleep-wake phenotypes for assessing maternal sleep health during pregnancy.

## Materials and Methods

### Animals

Adult male and female Wistar rats (3-5 months old) were obtained from Charles River Laboratories. Female rats were received naïve or timed pregnant on embryonic day (ED) 3. Upon arrival, all animals were singly housed with *ad libitum* food and water on a 12/12h light/dark cycle, where lights on corresponded to Zeitgeber time (ZT) 0 and lights off to ZT 12, in a temperature-controlled facility fully accredited by the American Association for the Accreditation of Laboratory Animal Care. All animals were allowed at least one week to acclimate to the animal facility before being used in experiments. All protocols were approved by the Institutional Animal Care and Use Committee at the University of South Carolina Floyd School of Medicine and were in accordance with the National Institutes of Health Guide for the Care and Use of Laboratory Animals.

### Chemicals

L-kynurenine sulfate salt (“kynurenine,” purity: 99.4%) was procured from Sai Advantium, Hyderabad, India. For liquid chromatography-tandem mass spectrometry (LC-MS/MS), unless specified otherwise, chemicals were obtained from Sigma Aldrich, St. Louis, Missouri, USA. All other reagents were obtained from various suppliers of the highest commercially available purity.

### Kynurenine Diet

#### Treatment in Pregnant Females

Pregnant females received kynurenine-supplemented diet, herein named embryonic kynurenine (EKyn), as described in previous publications [59], containing 100 mg kynurenine/day or control (ECon) wet mash daily for 8 days during the last week of gestation from ED 15 to ED 22. Each rat received approximately 30 g of wet mash daily to exceed anticipated daily requirements. Diet was administered between ZT 2 and ZT 4 each day. For biochemical analyses, dams (N = 7-8/group) were euthanized at ZT 6 on day seven of diet administration, ED 21, to harvest tissues before pregnant dams entered labor.

#### Treatment in Non-Pregnant Females

Adult female rats received kynurenine-supplemented diet containing 100 mg kynurenine/day or control wet mash daily for 8 days. Each rat received approximately 30 grams of wet mash per day. Diet was administered daily between ZT 2 and ZT 4. For biochemical analyses, females (N = 8-9/group) were euthanized at ZT 6 on day seven of diet administration to harvest tissues at an equivalent time point as ECon and EKyn dams.

### Sleep-Wake Monitoring

#### Surgical procedure

Females were implanted intraperitoneally with radiotelemetry devices (PhysioTel HD-S02, Data Sciences International (DSI), St. Paul, MN, USA) to acquire EEG and EMG for sleep monitoring as previously described [51,62]. Briefly, under isoflurane anesthesia, animals were placed in a stereotaxic frame (Stoelting Co., Wood Dale IL, USA) and carprofen (5 mg/kg, s.c.) was administered as an analgesic. EEG/EMG leads were threaded underneath the skin from a transverse incision into the abdominal cavity to reach a longitudinal incision along the midline of the head and neck. EEG leads were wrapped around two surgical screws positioned over the frontal and parietal lobes (2.0 mm anterior/+1.5 mm lateral and 7.0 mm posterior/-1.5 mm lateral relative to bregma) and secured with acrylic dental cement. EMG leads were sutured into the cervical neck muscle approximately 1.0mm apart. Incisions were closed with suture or wound clips. Animals were allowed one full week of recovery prior to sleep-wake data acquisition.

#### Sleep Monitoring in Pregnant Females

Adult female rats (N = 7 - 8/group) were implanted with EEG/EMG telemetry devices. Following at least one week of surgical recovery, sleep was continuously recorded during a 96-h undisturbed baseline period designed to span the female estrous cycle [64]. To allow for within-subject sleep-wake behavioral analysis, pregnancy was then induced by pairing females in proestrus overnight with a proven male breeder. A vaginal plug the next morning or a weight gain of at least 10 g after 3 days confirmed pregnancy. Females without a vaginal plug or significant weight gain 3 days post-pairing were re-paired with a different male the next time they entered proestrus. EKyn or ECon diet, described above, was administered from ED 15 to ED 22. At birth (postnatal day [PD] 0), females were returned to standard food. For analysis, sleep recordings were divided into three periods: i) baseline, consisting of the 96-h undisturbed pre-pregnancy recordings; ii) pregnancy, denoted ECon or EKyn as described above, consisting of 24-h recordings on ED 16, ED 18, and ED 20; and iii) postpartum, consisting of a 24-h recording on PD 5.

#### Sleep Monitoring in Non-Pregnant Females

Adult female rats (N = 8) were implanted with EEG/EMG telemetry devices. Following at least one week of surgical recovery, sleep was continuously recorded for i) baseline, 96-h undisturbed period designed to span the female estrous cycle; ii) kynurenine diet exposure, consisting of 24-h recordings on days 2, 4, and 6; and iii) washout, consisting of 24-h recording 5 days after return to standard food.

#### Sleep-wake data acquisition and analysis

Sleep data were continuously acquired at a sampling rate of 500 Hz on a PC with Ponemah 6.10 software (DSI). Digitized signal data were scored offline with an automated machine learning algorithm [65] and NeuroScore 3.4 (DSI) in 10-s epochs as wake (low-amplitude, high-frequency EEG and high-amplitude EMG), non-rapid eye movement sleep (NREM; high-amplitude, low-frequency EEG and little to no EMG activity), or rapid eye movement sleep (REM; low-amplitude, high-frequency EEG with little to no EMG activity). All recordings initially classified by long short-term memory (LSTM) were reviewed in NeuroScore by human experts. Data were analyzed in 12-h time bins for each vigilance state, assessing total duration, number of bouts, average bout duration, NREM and REM sleep onset, and the number of transitions between vigilance states. Discrete fast Fourier transform estimated the NREM and REM sleep power spectrum for each epoch in the delta (0.5–4 Hz), theta (4–8 Hz), alpha (8–12 Hz), sigma (12–16 Hz), and beta (16–20 Hz) frequency ranges, and normalized to the total power per subject. Relative home cage activity and core body temperature were reported by NeuroScore.

### Saliva Collection

Saliva was collected in both non-pregnant and pregnant rats fed kynurenine-supplemented diet, and respective controls. Saliva was obtained at the following time points each day: day 1 at ZT 6 and ZT 12; day 4 at ZT 0, ZT 6, and ZT 12; and day 7 at ZT 0 and ZT 6. Briefly, animals were lightly restrained, and a small cotton-tipped applicator was placed in the oral cavity until saturated with saliva (20-30s). This step was repeated such that two swabs were collected per subject to ensure ample saliva collection. Saliva-saturated applicators were placed in nested collection tubes comprised of a 0.5 mL microcentrifuge tube previously punctured at the bottom with a sterile 18G syringe needle and fit into an outer 1.5 mL microcentrifuge tube [66]. Collection tubes with applicators were immediately centrifuged for approximately 2 min (2000 g) to separate saliva (Thermo Scientific mySPIN 6 mini centrifuge). Approximately 10-50 μL of saliva could be collected total per subject. Saliva samples were snap frozen on dry ice and stored at -80°C.

### Tissue Collection

Animals were euthanized with CO_2_ asphyxiation, and whole trunk blood was collected in tubes containing 25 μL K3-EDTA (0.15%) as an anticoagulant. Blood was centrifuged (1000 g, 10 min) to separate plasma. Brains were promptly removed, dissected into regions of interest (brainstem, hypothalamus, hippocampus, frontal cortex) on wet ice, and snap-frozen on dry ice. All biological samples were stored at -80°C.

### Biochemical Analyses

#### Corticosterone

Corticosterone was analyzed in plasma (diluted 1:40 v/v with steroid displacement reagent) with an enzyme-linked immunosorbent assay (ELISA), according to manufacturer instructions (Enzo Life Sciences, Farmingdale, NY, USA). ELISA plates were read with a Synergy 2 Gen 5 microplate reader at 405 nm absorbance (BioTek Instruments Inc., Winooski, VT, USA).

#### Inflammatory Markers

Inflammatory markers were measured in an aliquot of maternal plasma (diluted 1:2 v/v in phospho-buffered saline, pH: 7.5) by a Rat Cytokine/Chemokine 27-Plex Discovery Assay Array (Eve Technologies, Calgary, AB, Canada) [22].

#### Tryptophan, Kynurenine, and Kynurenic Acid (KYNA)

The content of tryptophan, kynurenine, or KYNA was determined by ultra-high-performance liquid chromatography (UPLC). All samples were diluted in ultrapure water. Specific dilutions were as follows: maternal saliva (1:10 v/v for KYNA), maternal plasma (1:1000 v/v for tryptophan; 1:10 v/v for kynurenine; 1:10 v/v for KYNA), and maternal brain (sonicated 1:5 w/v, final dilution 1:10 for kynurenine and KYNA). Eighty μL of diluted sample was acidified with 20 mL of perchloric acid (6% saliva and plasma, 25% brain) and centrifuged (12,000 rpm, 10 min). Twenty or thirty μL of the supernatant was isocratically eluted from a ReproSil-Pur C18 column (4 mm x 100 mm; Dr. Maisch, GmbH, Ammerbuch, Germany), using a mobile phase containing 50 mM sodium acetate and 3-5% acetonitrile (pH adjusted to 6.2 with glacial acetic acid) at a flow rate of 0.5 mL/min, and detected in the eluate with 500 mM zinc acetate delivered after the column with a flow rate of 0.1 mL/min. In the eluate, tryptophan (excitation: 285 nm, emission: 365 nm), kynurenine (excitation: 365 emission: 480), and KYNA (excitation 344 nm, emission 398 nm) were detected fluorometrically (Acquity UPLC H-Class PLUS System, Waters Corporation, Bedford, MA, USA) with Empower 3 software (Waters Corporation). Brain tissue analysis was normalized to protein content using the Lowry method [67].

#### Quinolinic Acid (QUIN)

The content of QUIN in plasma was determined by liquid chromatography tandem mass spectrometry (LC-MS/MS) as previously described [22]. Briefly, a QUIN standard curve ranging from 5 fmol (limit of detection) to 50 pmol was prepared. 50 µL plasma was precipitated with ice-cold 80% methanol. 10 pmol QUIN-d3 internal standard (MedChemExpress, Monmouth Junction, NJ, USA), calibrated to the weight of freshly prepared QUIN, was added to each standard and sample. The standards and plasma samples were prepared for analysis by solid-phase extraction with 1cc C18 SPE cartridges (Waters Corporation) using 0.1% trifluoroacetic acid/40% methanol as the eluent before derivatization with 100:20 v/v ethanol/acetyl chloride, as described previously [68]. The reaction products were resuspended in 200 µL acetonitrile/water (20:80 v/v) and LC separation was performed on a Thermo Vanquish Flex liquid chromatography system (ThermoFisher Scientific) using a Waters XBridge C18 reverse-phase column. Following a 3 µL sample injection, a binary gradient elution was performed, with Solvent A consisting of water with 0.1% formic acid and solvent B containing acetonitrile with 0.1% formic acid. Positive ion electrospray mass spectra were acquired on a Thermo Q-Exactive HF-X Quadrupole-Orbitrap performing parallel reaction monitoring (PRM). XCalibur 4.2 software (ThermoFisher) was used to construct extracted ion chromatograms of the transitions 224→150 (QUIN), and 227→153 (QUIN-d3). The area of the QUIN peak was normalized to the area of the QUIN-d3 internal standard to obtain peak area ratios. The mass of QUIN in the plasma samples was normalized to the plasma volume to calculate QUIN concentration.

### Data Handling and Statistics

All data are presented as mean ± SEM. All statistical analyses were conducted utilizing GraphPad Prism 11.0 software (GraphPad Software, La Jolla, CA, USA), with a threshold for statistical significance set at *p* < 0.05. Average Salivary KYNA was calculated for each collection day. Plasma inflammatory markers were log_2_-transformed for heat map visualizations to highlight fold-changes between experimental groups and manage the wide range of concentrations. All statistical analyses were conducted on raw values (pg/mL). For biochemical data, outliers, as identified by a ROUT test with Q = 1%, were excluded from the analysis. Baseline sleep data represent the average of four days of sleep-wake behavior recording. Limited unforeseen system failures interrupted sleep data acquisition during periods of diet administration. Sleep data were evaluated in 12-h bins (ZT 0-12, light phase, and ZT 12-24, dark phase) with all post hoc comparisons conducted against the non-pregnant baseline. Whenever possible, a Geisser-Greenhouse correction was applied to ANOVA tests. All significant two-way ANOVA results were subsequently analyzed using Fisher’s LSD post hoc test. A supplementary statistical file has been provided to present all ANOVA results.

Salivary KYNA was evaluated using a two-way repeated measures (RM) ANOVA, with treatment (control or kynurenine) as a between-subject factor and treatment day as a within-subject factor. Non-pregnant and pregnant females were assessed separately. Spearman correlational analyses were used to examine the relationship between average salivary KYNA levels and brain KYNA levels. Remaining biochemical data from non-pregnant control and non-pregnant kynurenine females were analyzed with Mann-Whitney tests. To examine the effect of pregnancy, biochemical data were analyzed using two-way ANOVA with post hoc comparisons within control or kynurenine treatment conditions. Each plasma inflammatory marker and brain region was analyzed independently.

In pregnant females, NREM and REM sleep at baseline, ED 16, ED 18, and ED 20 were evaluated using two-way RM ANOVA with pregnancy status and light phase as within-subject factors. The change from non-pregnant baseline was calculated for postpartum sleep and compared to baseline with Mann-Whitney tests. Comparisons between treatment groups were made by two-way ANOVA with treatment group as a between-subject factor and light phase as a within-subject factor. Remaining sleep-wake data were evaluated for all treatment days using two-way RM ANOVA, with treatment day and light phase as within-subject factors. Spectral power was assessed by two-way RM ANOVA with frequency and pregnancy status as within-subject factors.

## Results

### Salivary KYNA can distinguish control from kynurenine-fed female rats

We administered control or kynurenine-supplemented diet to pregnant (**Figure 1A**) and non-pregnant (**Figure 1B**) Wistar rats. During pregnancy, embryonic control (ECon) or embryonic kynurenine (EKyn) diets were administered during the last week of gestation from embryonic day (ED) 15 to ED 22, a period highly sensitive to prenatal and KP perturbations [22,23,59]. In non-pregnant females, diet was administered for eight consecutive days to align with ECon and EKyn treatments. In pregnant females, kynurenine diet (EKyn) does not influence plasma inflammatory markers, yet elevates KP metabolism and brain KYNA compared to ECon [22]. Clinical studies have investigated salivary KYNA as a potential biomarker for brain health [69-71]. Our goal was to establish a method for rat salivary KYNA determination and test whether salivary KYNA could distinguish kynurenine-fed from control-fed females. Importantly, we detected KYNA in the nanomolar range across all collected samples. In pregnant females, EKyn elevated salivary KYNA compared to ECon (main effect EKyn: *F*_1, 12_ = 7.301, *p* < 0.05; ED 21: *p* < 0.05) (**Figure 1C**). In non-pregnant females, salivary KYNA was similarly elevated in kynurenine-fed females compared to control (main effect kynurenine: *F*_1, 15_ = 17.18; *p* < 0.001; day 1: *p* < 0.05, day 4: *p* < 0.01, day 6: *p* < 0.05) (**Figure 1D**). Positive correlations were observed between salivary and brain KYNA in the brainstem (*r*_s_ = 0.5248, *p* < 0.01), hypothalamus (*r*_s_ = 0.5378, *p* < 0.01), cortex (*r*_s_ = 0.5623, *p* < 0.01), hippocampus (*r*_s_ = 0.3637, *p* < 0.05), and frontal cortex (*r*_s_ = 0.504, *p* < 0.05) (**Figure S1**). We confirmed that the kynurenine diet in non-pregnant females did not affect plasma corticosterone or tryptophan; however, it predictably increased plasma kynurenine, KYNA, and QUIN, indicating enhanced peripheral kynurenine pathway metabolism. Importantly, kynurenine diet elevated KYNA levels throughout the female brain. (**Figure 2**). Kynurenine diet did not influence plasma inflammatory markers in non-pregnant females (**Figure S3**). We also report the influence of pregnancy on KP metabolite levels in plasma (**Figure S4**) and brain (**Figure S5**). Regardless of kynurenine diet status, pregnancy increased peripheral KP metabolism from tryptophan and KYNA levels in the brainstem, hippocampus, and frontal cortex.

**Figure 1:**
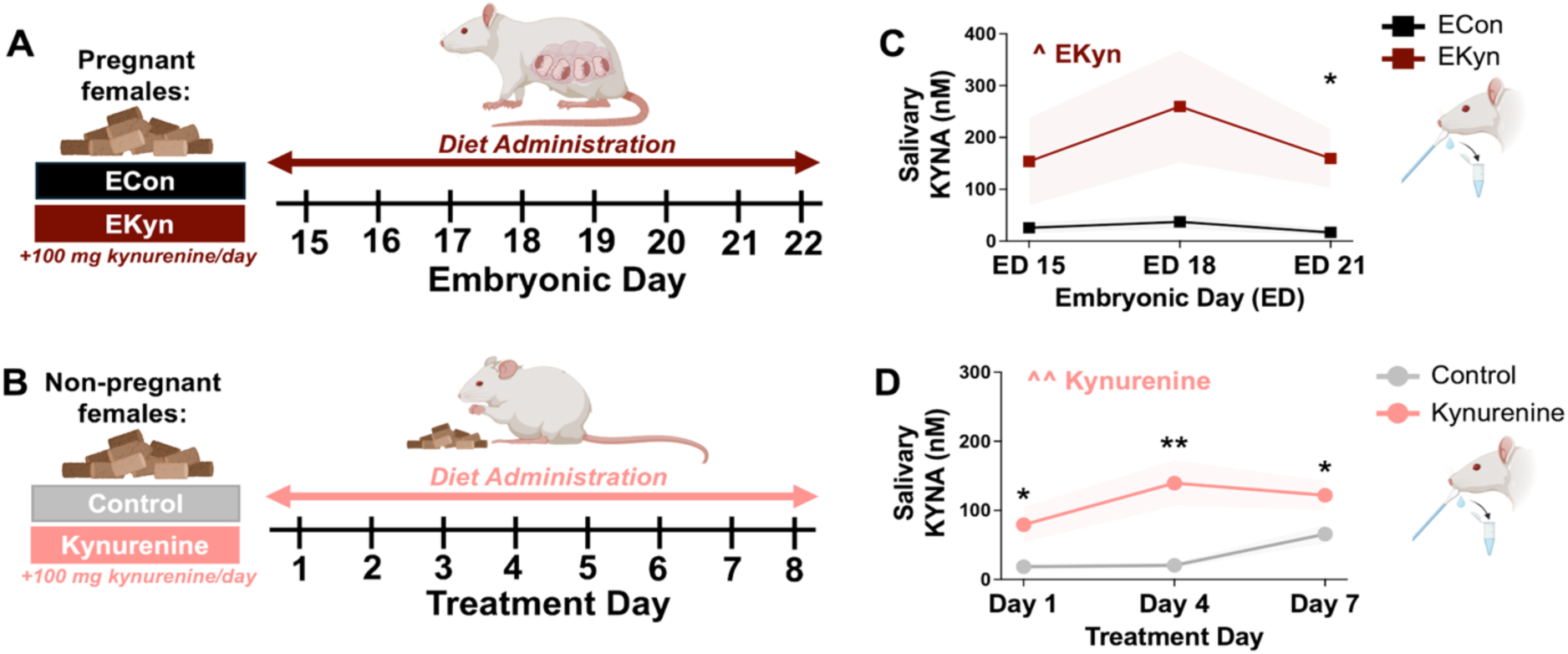
Salivary KYNA distinguishes kynurenine-fed females from control-fed females. **(A)** Pregnant females received control (Embryonic Control, ECon) or kynurenine-supplemented diet (Embryonic Kynurenine, EKyn, + 100 mg/day kynurenine) for 8 days from embryonic day (ED) 15 to ED 22. **(B)** Non-pregnant females received control diet or kynurenine-supplemented diet (+ 100 mg/day kynurenine) for 8 days. **(C)** Salivary KYNA levels in pregnant females. **(D)** Salivary KYNA levels in non-pregnant females. Data are mean ± SEM. Two-way RM ANOVA: ^ *p* < 0.05. Fisher’s LSD post-hoc: * *p* < 0.05, ** *p* < 0.01. N = 6-9/group.

**Figure 2:**
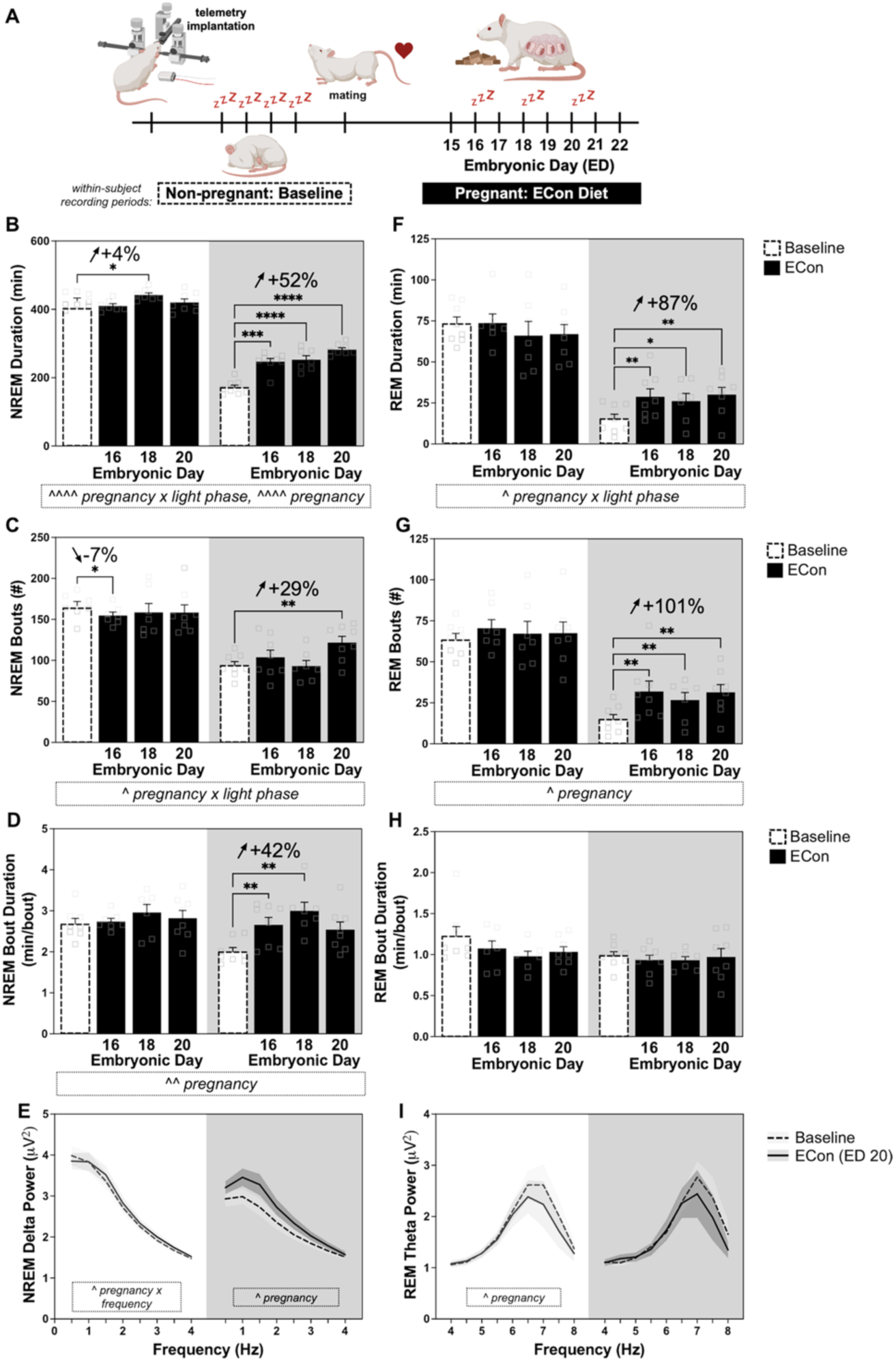
Pregnancy significantly increases dark phase NREM and REM sleep duration and alters dark phase sleep architecture in ECon dams. **(A)** Experimental timeline. Sleep was recorded during a non-pregnant baseline and on embryonic day (ED) 16, ED 18, and ED 20. Females received embryonic control diet (ECon) from ED 15 to ED 22. Data are separated by light phase, shown with a white background, and dark phase, shown with a grey background. NREM sleep outcomes include **(B)** duration, **(C)** bout number, **(D)** bout duration, and **(E)** delta power. REM sleep outcomes include **(F)** duration, **(G)** bout number, **(H)** bout duration, and **(I)** theta power. Data are mean ± SEM. Two-way RM ANOVA. Significant interactions and main effects of ECon are displayed in boxed annotations below each figure panel: ^ *p* < 0.05, ^^ *p* < 0.01, ^^^^ *p* < 0.0001. Fisher’s LSD post-hoc: * *p* < 0.05, ** *p* < 0.01, **** *p* < 0.0001. N = 6-8/day.

### Pregnancy significantly increases dark phase sleep in the last week of gestation

We have observed limited changes to maternal sleep-wake architecture between ECon and EKyn dams when compared only at ED 20, immediately prior to birth [22]. However, pregnancy-associated changes in sleep-wake behavior and intrinsic variability in rodent sleep patterns may obscure treatment-related effects in between-group comparisons. Therefore, we adopted a within-subjects design, comparing each treatment day with the corresponding animal’s own non-pregnant baseline. We verified that baseline sleep-wake architecture did not differ significantly between experimental cohorts (**Figure S6**).

To characterize maternal sleep during the last week of gestation, we recorded control rats during a non-pregnant baseline period before mating and again during 24-h periods on ED 16, ED 18, and ED 20. During this gestational period, the embryonic control diet, ECon, was administered daily (**Figure 2A**). When we assessed NREM sleep, we found a significant pregnancy x light phase interaction (*F*_1.809, 10.85_ = 30.72, *p* < 0.0001) and a main effect of pregnancy (*F*_2.547, 17.83_ = 19, *p* < 0.0001) (**Figure 2B**). Most notably, dark phase NREM sleep increased by 52% compared to non-pregnant baseline across all embryonic days (ED 16: *p* < 0.001, ED 18: *p* < 0.0001, ED 20: *p* < 0.0001). NREM bout number was also affected by a significant pregnancy x light phase interaction (*F*_1.948, 11.69_ = 5.663, *p* < 0.05), with pregnancy decreasing NREM bouts during the light phase at ED 16 (-7%, *p* < 0.05) and increasing NREM bouts during the dark phase at ED 20 (+29%, *p* < 0.05) (**Figure 2C**). Pregnancy also significantly affected NREM bout durations (main effect of pregnancy: *F*_1.664, 11.65_ = 8.026, *p* < 0.01) (**Figure 2D**). During the dark phase, NREM bout duration increased by an average of 42% at ED 16 (*p* < 0.01) and ED 18 (*p* < 0.01) compared with non-pregnant baseline.

To explore the effects of pregnancy on NREM sleep quality and sleep pressure, we calculated NREM delta power on ED 20, when dams were near parturition. Pregnancy altered light phase NREM delta power (pregnancy x frequency interaction: *F*_1, 4.857_ = 7.778, *p* < 0.05). During the dark phase, a main effect of pregnancy (*F*_1, 6_ = 9.916, *p* < 0.05) revealed increased NREM delta power compared to non-pregnant baseline (1 Hz: *p* < 0.05, 1.5 Hz: *p* < 0.05, 2.5 Hz: *p* < 0.05, 3 Hz: *p* < 0.05) (**Figure 2E**). Together, these findings indicate that late pregnancy increases dark phase NREM sleep and alters NREM sleep architecture, consistent with heightened sleep pressure near parturition.

Pregnancy also significantly impacted REM sleep across the last week of gestation. A pregnancy x light phase interaction (*F*_1.789, 10.73_ = 6.548, *p* < 0.05) affected total REM sleep duration (**Figure 2F**). During the dark phase, REM sleep duration increased 96% on ED 16 (*p* < 0.01) and ED 20 (*p* < 0.01) compared with non-pregnant baseline. This increase was driven primarily by increased dark phase REM sleep bout number (**Figure 2G**, main effect of pregnancy: *F*_1.767, 19.37_ = 5.065, *p* < 0.05), which was elevated across embryonic days (ED 16: *p* < 0.01, ED 18: *p* < 0.01, ED 20: *p* < 0.01). In contrast, REM sleep bout duration was not impacted by pregnancy (**Figure 2H**). We next calculated REM theta power, the dominant frequency band during REM sleep, on ED 20. Pregnancy reduced light phase REM theta power compared with baseline (main effect pregnancy: *F*_1, 6_ = 9.596, *p* < 0.05), with significant reductions at 7 Hz and 7.5 Hz (both *p* < 0.05) (**Figure 2I**). REM theta power did not change during the dark phase. Together, these findings indicate that late pregnancy increases dark phase REM sleep primarily through more frequent REM episodes, while selectively reducing light phase REM theta power near parturition.

### Delayed impairment to REM sleep in EKyn dams

Next, we characterized the effects of kynurenine exposure during pregnancy by comparing sleep-wake architecture in EKyn dams on ED 16, ED 18, and ED 20 to each animal’s own non-pregnant baseline. EKyn diet was administered from ED 15 to ED 22 (**Figure 3A**). During pregnancy, EKyn dams showed a significant pregnancy x light phase interaction (*F*_1.628, 8.683_ = 10.95, *p* < 0.01) and a main effect of pregnancy (*F*_2.107, 12.64_ = 11.54, *p* < 0.01) for NREM duration (**Figure 3B**). Like ECon dams, EKyn dams showed a 52% increase in dark phase NREM sleep across ED 16 (*p* < 0.001), ED 18 (*p* < 0.0001), and ED 20 (*p* < 0.01) compared with non-pregnant baseline. A pregnancy x light phase interaction (*F*_1.737, 9.266_ = 7.148, *p* < 0.05) also influenced NREM bout number (**Figure 3C**). On ED 16, EKyn dams had fewer NREM bouts during the light phase (-12%, *p* < 0.05) and more NREM bouts during the dark phase (+13%, *p* < 0.05) compared with their non-pregnant baseline, whereas NREM bout number was unaffected on ED 18 or ED 20. NREM bout duration was also influenced by a significant pregnancy x light phase interaction (*F*_1.83, 9.761_ = 11.63, *p* < 0.01) and a main effect of pregnancy (*F*_2.125, 12.75_ = 3.936, *p* < 0.05) (**Figure 3D**), with dark phase NREM bout duration increased 38% across ED 16 (*p* < 0.05), ED 18 (*p* < 0.01), and ED 20 (*p* < 0.01). Finally, pregnancy (interaction: *F*_1.843, 8.949_ = 6.609, *p* < 0.05; main effect pregnancy: *F*_1, 6_ = 28.15, *p* < 0.01) increased dark phase NREM delta power in EKyn dams (1 Hz: *p* < 0.01, 1.5 Hz: *p* < 0.01, 2 Hz: *p* < 0.001, 2.5 Hz: *p* < 0.05, 3 Hz: *p* < 0.01, 3.5 Hz: *p* < 0.05) (**Figure 3E**).

**Figure 3:**
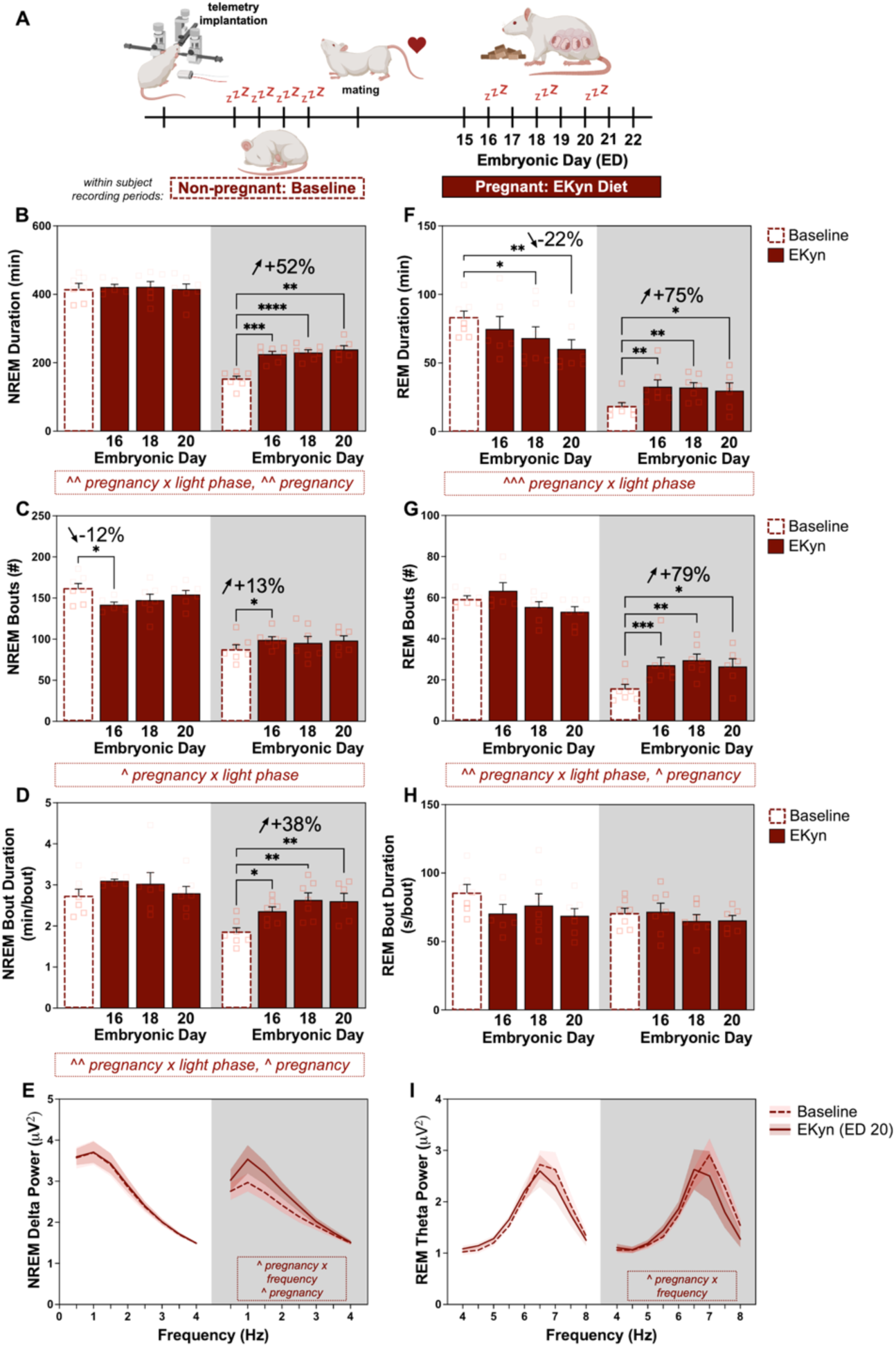
EKyn preserves NREM sleep adaptations to pregnancy but reduces light phase REM sleep beginning on ED 18. **(A)** Experimental timeline. Sleep was recorded during a non-pregnant baseline and on embryonic day (ED) 16, 18, and 20. Females received embryonic kynurenine diet (EKyn; + 100 mg/day kynurenine) from ED 15 to ED 22. Data are separated by light phase, shown with a white background, and dark phase, shown with a grey background. NREM sleep outcomes include **(B)** duration, **(C)** bout number, and **(D)** bout duration. REM sleep outcomes include **(E)** duration, **(F)** bout number, and **(G)** bout duration. Data are mean ± SEM. Two-way RM ANOVA. Significant interactions and main effects of EKyn are displayed in boxed annotations below each figure panel: ^ *p* < 0.05, ^^ *p* < 0.01, ^^^ *p* < 0.001. Fisher’s LSD post-hoc: * *p* < 0.05, ** *p* < 0.01, *** *p* < 0.001, **** *p* < 0.0001. N = 6-7/day.

Analysis of REM sleep duration in EKyn dams revealed a significant pregnancy x light phase interaction (*F*_1.871, 9.987_ = 16.32, *p* < 0.001) (**Figure 3F**). In contrast to ECon dams, EKyn dams showed reduced light phase REM sleep beginning at ED 18, with REM duration decreasing by 22% at ED 18 (*p* < 0.05) and ED 20 (*p* < 0.01) compared with non-pregnant baseline. However, EKyn dams retained pregnancy-associated increases in dark phase REM sleep at ED 16 (*p* < 0.01), ED 18 (*p* < 0.05), and ED 20 (*p* < 0.05). REM bout number was influenced by a significant pregnancy x light phase interaction (*F*_1.744, 10.303_ = 8.574, p < 0.01) and a main effect of pregnancy (*F*_1.674, 10.05_ = 5.013, *p* < 0.05) (**Figure 3G**). Specifically, dark phase REM bout number increased by 79% across all embryonic days assessed (ED 16: *p* < 0.001, ED 18: *p* < 0.01, ED 20: *p* < 0.05). Pregnancy did not significantly affect REM bout duration, although the pregnancy x light phase interaction approached statistical significance (*F*_1.922, 10.25_ = 3.418, *p* = 0.074) (**Figure 3H**). Finally, dark phase REM theta power was influenced by a significant pregnancy x frequency interaction (*F*_1.658, 8.082_ = 4.758, *p* < 0.05) (**Figure 3I**), with theta power significantly reduced in EKyn dams compared to their non-pregnant baseline (5.5 Hz: p < 0.05, 7.5 Hz: *p* < 0.01, 8 Hz: *p* < 0.01).

### In ECon, postpartum sleep timing is altered, and wakefulness is fragmented

In the postpartum period, disrupted maternal sleep-wake patterns often persist or emerge due to maternal caregiving demands [72]. These disruptions are also associated with increased risks of adverse maternal and offspring health outcomes [73-75]. As such, we characterized the postpartum sleep architecture in controls on PD 5, providing mothers time to acclimate after birth. When we assessed sleep duration, light phase NREM sleep was significantly reduced compared with baseline (*p* < 0.05), while dark phase NREM sleep (*p* < 0.05) and REM sleep (*p* < 0.05) were increased (**Figure 4A**). Although light phase REM sleep duration was unchanged postpartum, REM bout number was reduced compared with baseline (*p* < 0.05). In contrast, dark phase NREM and REM bout numbers were increased postpartum (NREM: *p* < 0.05, REM: *p* < 0.05) (**Figure 4B**), consistent with greater fragmented sleep during the active phase. NREM bout duration was also reduced in the light phase (*p* < 0.05) (**Figure 4C**).

**Figure 4:**
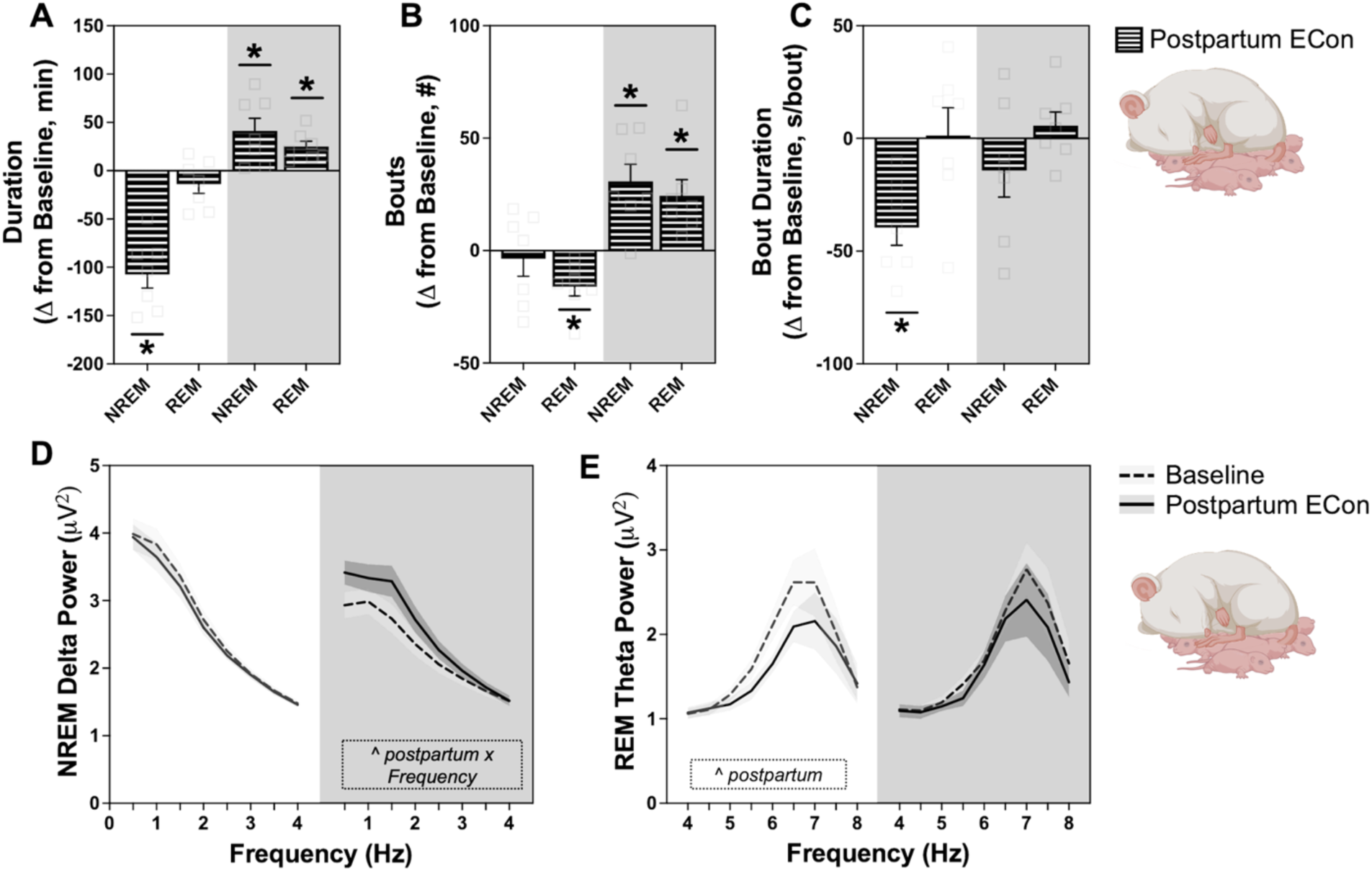
Postpartum disrupts sleep architecture in ECon dams. Data are separated by light phase, shown with a white background, and dark phase, shown with a grey background. Sleep outcomes are shown as change from non-pregnant baseline and include **(A)** vigilance-state duration, **(B)** bout number, and **(C)** bout duration. Spectral outcomes include **(D)** NREM sleep delta power and **(E)** REM sleep theta power. Data are mean ± SEM. Two-way RM ANOVA. Significant main effects are displayed in boxed annotations within each figure panel: ^ *p* < 0.05. Wilcoxon test: * *p* < 0.05. N = 6-7/day.

Regarding spectral power, dark phase NREM delta power was affected by a significant postpartum and frequency interaction (*F*_0.931, 4.522_ = 7.398, *p* < 0.05) and a main effect of postpartum status (*F*_1, 6_ = 25.33, *p* < 0.01). Compared with baseline, dark phase NREM delta power was increased (1.5 Hz: *p* < 0.05, 2 Hz: *p* < 0.01, 2.5 Hz, *p* < 0.001, 3 Hz: *p* < 0.05) (**Figure 4D**). REM theta power was also altered postpartum, with a main effect of postpartum status during the light phase (*F*_1, 6_ = 12.60, *p* < 0.05). Specifically, light phase REM theta power was reduced compared with baseline (5 Hz: *p* < 0.01, 5.5 Hz: *p* < 0.01, 6 Hz: *p* < 0.05, 6.5 Hz: *p* < 0.05) (**Figure 4E**). Changes in maternal wakefulness, relative cage activity, core body temperature (**Figure S7**), and sleep-wake transitions (**Table S1**) during pregnancy and postpartum are described in the supplemental materials.

### Postpartum sleep alterations in EKyn dams

In the postpartum period, EKyn dams showed reduced light phase NREM sleep (*p* < 0.05) and increased dark phase NREM (*p* < 0.05) and REM sleep (p < 0.05) (**Figure 5A**). REM bout number was reduced during the light phase (*p* < 0.05), whereas NREM and REM bout numbers were increased during the dark phase (NREM: *p* < 0.05, REM: *p* < 0.05) (**Figure 5B**). NREM bout duration was selectively reduced during the light phase (*p* < 0.05) (**Figure 5C**).

**Figure 5:**
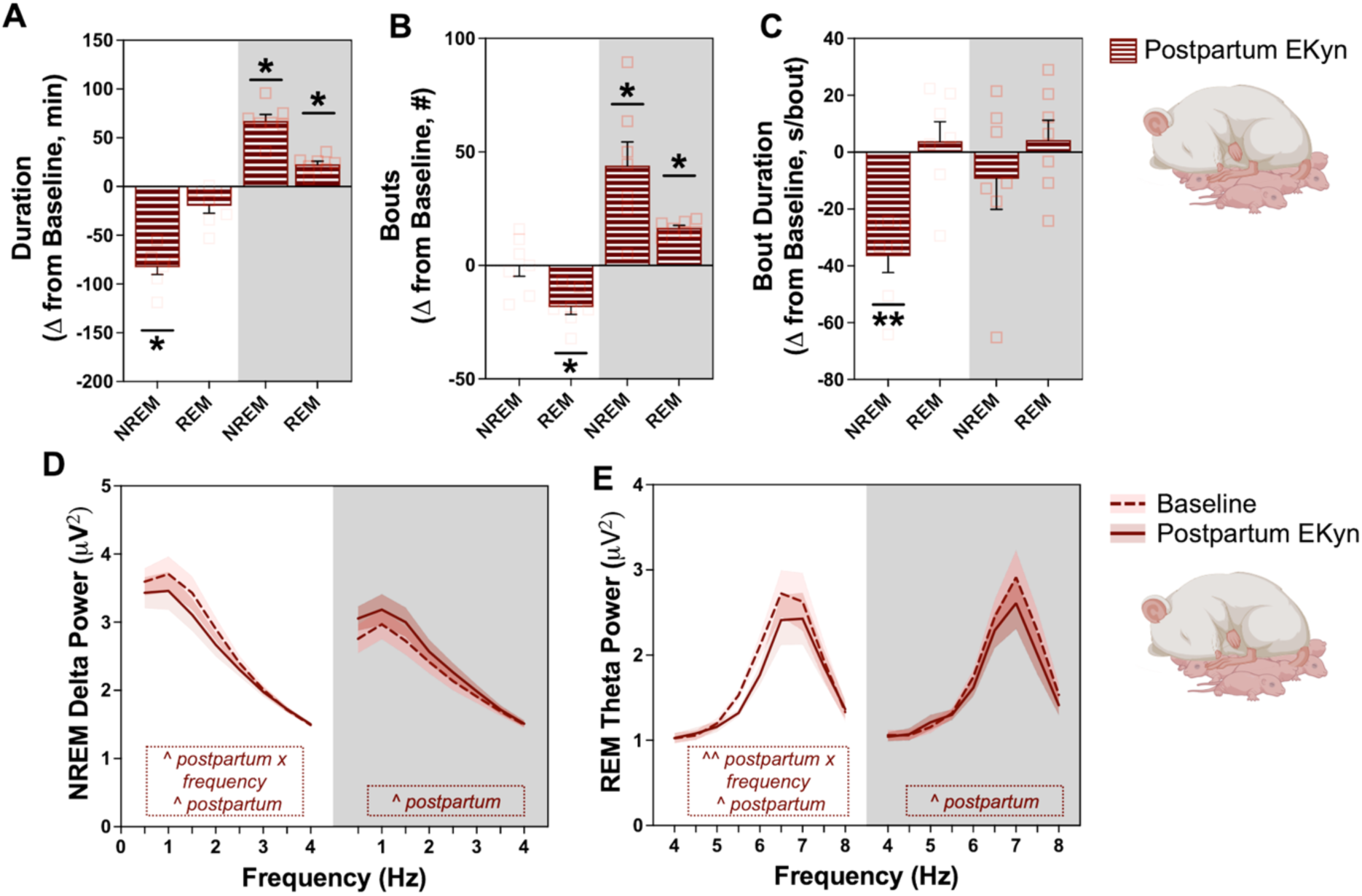
EKyn dams exhibit altered sleep architecture postpartum. Data are separated by light phase, shown with a white background, and dark phase, shown with a grey background. NREM and REM sleep outcomes are shown as change from non-pregnant baseline and include **(A)** vigilance-state duration, **(B)** bout number, and **(C)** bout duration. Spectral outcomes include **(D)** NREM sleep delta power and **(E)** REM sleep theta power. Data are mean ± SEM. Two-way RM ANOVA. Significant main effects are displayed in boxed annotations within each figure panel: ^ *p* < 0.05, ^^ *p* < 0.01. Wilcoxon test: * *p* < 0.05, ** *p* < 0.01. N = 7/group.

Spectral analyses revealed that light phase NREM delta power was affected by a significant postpartum x frequency interaction (*F*_1.671, 10.03_ = 4.554, *p* < 0.05) and a main effect of postpartum status (*F*_1, 6_ = 7.690, *p* < 0.05) at 1.5 Hz (*p* < 0.05) and 2 Hz (*p* < 0.01) (**Figure 5D**). In contrast, dark phase NREM delta power was increased postpartum, with a significant main effect of postpartum status (*F*_1, 6_ = 6.167, *p* < 0.05) and significant increases at 1 Hz (*p* < 0.05) and 1.5 Hz (*p* < 0.05). REM theta power was significantly reduced in postpartum EKyn dams during both the light phase (main effect postpartum status: *F*_1, 6_ = 6.167, *p* < 0.05; 5.5 Hz: *p* < 0.01, 6 Hz: *p* < 0.01) and dark phase (main effect postpartum status: *F*_1, 6_ = 7.227, *p* < 0.05; 7 Hz: *p* < 0.05, 7.5 Hz: *p* < 0.05, 8 Hz: *p* < 0.05) (**Figure 5E**).

Changes in maternal wakefulness, relative cage activity, core body temperature (**Figure S8**), and sleep-wake transitions (**Table S2**) during pregnancy and postpartum are described in the supplemental materials. We also compared ECon and EKyn sleep during pregnancy (**Figure S9**) and postpartum (**Figure S10**) and found no significant differences between groups during either time period.

### Kynurenine diet in non-pregnant females reduces light phase REM sleep

The effects of the kynurenine diet during pregnancy (EKyn) were delayed and less pronounced than expected based on prior kynurenine challenge studies in non-pregnant female rats, in which peripheral kynurenine injections acutely elevated brain KYNA levels and rapidly reduced REM sleep duration [62]. In contrast, dams consuming elevated kynurenine daily did not exhibit REM sleep deficits until the fourth day of diet administration. Because chronic KP activation and elevated brain KYNA are clinically relevant in females independent of pregnancy status, we included non-pregnant females both as a control for pregnancy-related effects and as a translationally relevant model for evaluating the impact of chronic kynurenine exposure on female sleep.

To maintain a within-subject design in non-pregnant females, sleep was recorded at baseline and during 24-h periods on days 2, 4, and 6 of kynurenine diet administration. A 24-h washout period was also recorded five days after cessation of the diet (**Figure 6A**). We first assessed NREM sleep. A significant treatment day x light phase interaction influenced NREM sleep duration (*F*_2.462, 14.16_ = 4.457, *p* < 0.05). Light phase NREM sleep was unaffected by kynurenine diet; however, during the dark phase, NREM sleep increased by an average of 28% on days 4 (*p* < 0.05) and 6 (*p* < 0.01), as well as during washout (*p* < 0.05) (see **Figure 6B**). Kynurenine diet did not significantly affect NREM bout number (**Figure 6C**) or NREM bout duration (**Figure 6D**).

**Figure 6:**
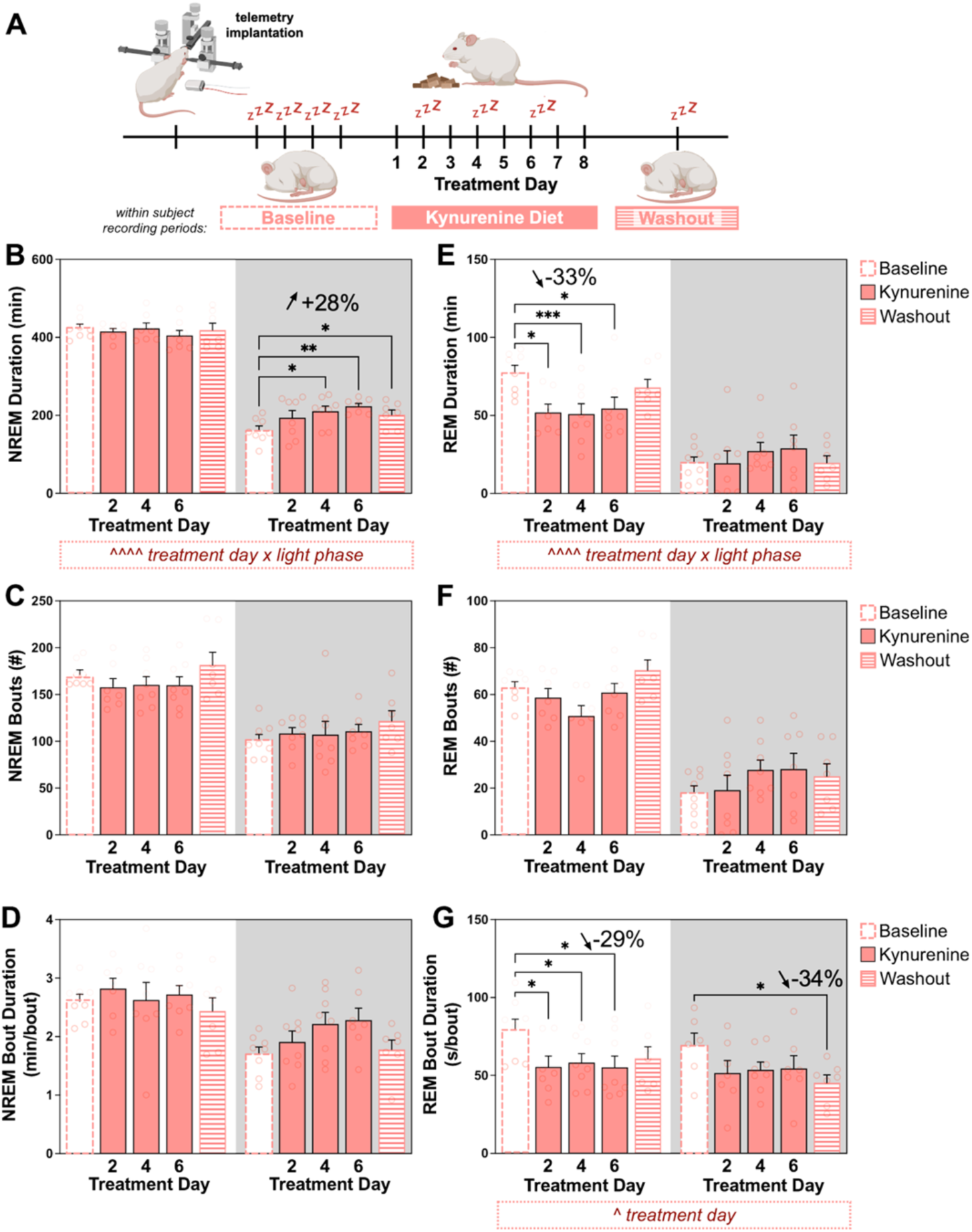
Kynurenine diet reduces light phase REM sleep and increases dark phase NREM sleep in non-pregnant females. **(A)** Experimental timeline. Sleep was recorded at baseline and on days 2, 4, and 6 during diet administration. Females received kynurenine-supplemented diet (+ 100 mg/day kynurenine) for 8 days. Data are separated by light phase, shown with a white background, and dark phase, shown with a grey background. NREM sleep outcomes include **(B)** duration, **(C)** bout number, and **(D)** bout duration. REM sleep outcomes include **(E)** duration, **(F)** bout number, and **(G)** bout duration. Data are mean ± SEM. Two-way RM ANOVA. Significant interactions and main effects of treatment day are displayed in boxed annotations below each figure panel: ^ *p* < 0.05, ^^^^ *p* < 0.0001. Fisher’s LSD post-hoc: * *p* < 0.05, ** *p* < 0.01, *** *p* < 0.001. N = 7-8/day.

**Figure 7:**
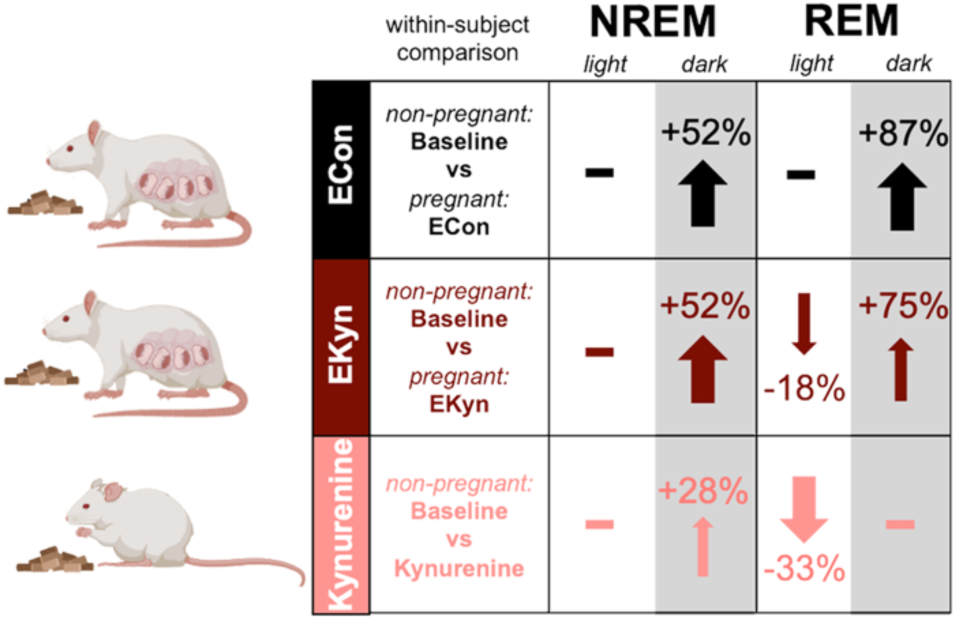
Pregnancy reduces the effects of kynurenine diet on female sleep. Pregnancy increases total sleep during the dark phase. EKyn dams retain much of this adaptation, yet experience reduced light phase REM sleep, an effect induced to a greater extent by kynurenine diet in non-pregnant females. Our results reveal a distinct role for chronic kynurenine exposure in female sleep regulation during pregnancy.

Strikingly, kynurenine diet reduced light phase REM sleep durations by 33% across treatment days (day 2: *p* < 0.05 day 4: *p* < 0.001, day 6: *p* < 0.05) while leaving dark phase REM sleep unchanged, as indicated by a significant treatment day x light phase interaction (*F*_1.377, 8.264_ = 5.844, *p* < 0.05) (**Figure 6E**). This light phase REM deficit was transient, as REM sleep duration returned to baseline levels after cessation of the diet. Changes in REM sleep duration were not attributable to altered REM bout number (**Figure 6F**) but instead reflected reduced REM bout duration across treatment days (main effect treatment day: *F*_1.88, 13.16_ = 4.877, *p* < 0.05; day 2: *p* < 0.05, day 4: *p* < 0.05, day 6: *p* < 0.05) (**Figure 6G**). REM bout duration was also reduced during the dark phase at washout (*p* < 0.05). Kynurenine diet did not alter NREM delta power or REM theta spectral power (**Figure S11).**

Changes in wakefulness, relative cage activity, and core body temperature are described in the supplemental materials (**Figure S12**). Kynurenine diet did not alter sleep-wake transitions (**Table S3)** or NREM and REM latencies across treatment periods (**Table S4**).

## Discussion

Our study aimed to characterize the impacts of pregnancy and chronic KP activation on maternal sleep during the final week of gestation in rats. To stimulate KP metabolism, rats were maintained on a diet supplemented with kynurenine, the initial KP metabolite. Pregnancy induced a dark phase-dependent increase in maternal sleep, a pattern largely maintained in kynurenine-fed EKyn dams. However, near parturition, EKyn dams exhibited a decline in light phase REM sleep that was not observed in pregnant controls. In non-pregnant females, kynurenine diet consistently diminished light phase REM sleep. Notably, in EKyn dams, REM sleep deficits were detectable only when compared with each animal’s own non-pregnant sleep-wake baseline, rather than with pregnant controls. This finding underscores the importance of individual sleep-wake physiology when evaluating sleep disturbances during dynamic physiological periods such as pregnancy. Collectively, these findings demonstrate that pregnancy alters the effects of enhanced KP metabolism and elevated brain KYNA in sleep-wake homeostasis during gestation and highlight the value of individual sleep-wake phenotypes for assessing maternal sleep health.

### Salivary KYNA as a translational biomarker of kynurenine exposure

We successfully detected salivary KYNA in rats, enabling differentiation between control- and kynurenine-treated groups. Salivary KYNA positively correlated with brain KYNA across numerous brain regions. In human studies, increased salivary KYNA has been linked to stress responsivity and central nervous system function in schizophrenia [69-71]. From a clinical perspective, salivary KYNA holds promise as a non-invasive peripheral biological marker that may help identify or predict symptoms associated with increased brain KYNA, including cognitive and sleep disturbances. To the best of our knowledge, this is the first study to quantify salivary KYNA in rats, demonstrating the feasibility of using rat models to assess salivary KYNA as a peripheral biomarker of central nervous system KYNA. Moreover, the ability to detect treatment-related increases in salivary KYNA in both pregnant and non-pregnant rats after extended kynurenine exposure, along with positive correlations between salivary and brain KYNA across these cohorts, represents an important initial step toward evaluating the utility of salivary KYNA as a translational peripheral biomarker for brain KYNA status.

### Pregnancy-associated sleep remodeling in rats

Pregnant controls demonstrated increased dark phase NREM and REM sleep during the final week of gestation compared with non-pregnant baseline. These increases were attributable to longer NREM sleep bouts and more frequent NREM and REM bouts. In humans, late pregnancy is often characterized by nighttime sleep fragmentation, reduced sleep quality, and increased daytime sleepiness, with daytime napping commonly reported as a compensatory sleep behavior [76-79]. In this context, the increased dark phase NREM sleep observed in pregnant rats may reflect an analogous increase in sleep opportunity during the active phase, supported by both longer and more frequent NREM bouts. Although human studies often report reduced or disrupted nighttime sleep during late pregnancy, our findings align with existing rodent studies showing increased sleep during the normally active phase of late gestation [13,14,16]. Species differences may reflect several factors, including reduced effects of fetal positioning on maternal sleep disruption, absence of nocturia, and unrestricted opportunity to sleep throughout the 24-h period in rodents versus humans [80,81]. Nonetheless, both humans and rodents appear to experience heightened sleep near the end of pregnancy, which we observed as increased dark phase NREM delta power, a marker of sleep pressure [82]. Increased sleep need during late gestation may support the metabolic demands of fetal growth and development prior to birth [83-85]. Furthermore, hormonal fluctuations, including elevated levels of progesterone and its neuroactive metabolite allopregnanolone, prolactin, and estrogen, are widely recognized as key neurobiological factors that contribute to maternal sleep alterations in both humans and rodents [13,14,16,86,87]. These findings highlight analogous physiological processes in humans and rats during pregnancy and support the translational value of rodent models for studying pregnant sleep.

### Pregnancy attenuates but does not eliminate kynurenine-associated REM sleep disruption

By adopting a within-subject design, we detected kynurenine-induced sleep impairments during pregnancy that were not apparent in previous studies assessing sleep only at ED 20 [22]. Although EKyn dams retained a pregnancy-associated increase in dark phase NREM and REM sleep, they progressively lost light phase REM sleep with continued diet administration and advancing gestation. This reduction may be particularly consequential during pregnancy, as REM sleep is closely linked to synaptic plasticity and memory consolidation [88], and the maternal brain undergoes significant remodeling across cortical, limbic, and sensory systems throughout gestation [24,89-94]. Thus, reduced REM sleep near the end of pregnancy may represent a biologically meaningful disruption during a period of heightened maternal neural plasticity. REM-related alterations in hippocampal or amygdala function could contribute to maternal cognitive difficulties and increased vulnerability to poor mental health during the perinatal period [93,94].

In pregnant rats fed a kynurenine-supplemented diet (EKyn), the delayed emergence of REM sleep loss was unexpected, as acute peripheral kynurenine administration rapidly reduces REM sleep in non-pregnant females [62]. Differences in exposure route, timing, and treatment duration may partially explain the divergent effects of kynurenine challenge and chronic dietary kynurenine supplementation. However, pregnancy-associated physiological adaptations likely also contribute to this delayed response. Pregnancy is accompanied by extensive neuroendocrine, metabolic, and neural adaptations that support maternal physiology and fetal development, many of which also influence sleep regulation and responses to physiological challenge [95,96]. These adaptations may initially reduce the sensitivity of maternal sleep-wake circuits to elevated kynurenine metabolism, delaying the emergence of REM sleep deficits despite increased brain KYNA. This interpretation is supported by the immediate and persistent reductions in light phase REM sleep observed in non-pregnant females fed a kynurenine-supplemented diet. Thus, pregnancy may transiently buffer kynurenine-associated sleep disruption before susceptibility emerges later in gestation. To our knowledge, these findings provide the first evidence that pregnancy modulates the effects of kynurenine on sleep. Significantly, REM sleep deficits induced by kynurenine in non-pregnant females persisted across treatment days; however, they were reversible following a washout period. This observation suggests that sleep impairments may endure while KP activation remains elevated and subsequently recover once KP metabolism returns to normal.

### Brain KYNA as a mediator of kynurenine-associated sleep disruption

Regardless of pregnancy status, we propose that kynurenine disrupts sleep by increasing brain KYNA, a downstream KP metabolite. Kynurenine is transported from plasma into the brain and subsequently converted to KYNA in astrocytes via kynurenine aminotransferases (KATs) [97]. Newly synthesized KYNA is rapidly released into the extracellular space, where it exerts neuromodulatory effects. Preclinical studies show that elevated brain KYNA levels impair sleep-wake behavior, whereas reduced brain KYNA improves sleep outcomes, supporting a role for KYNA in sleep regulation [50,51,62,63]. Peripheral kynurenine administration rapidly increases brain KYNA and produces transient REM sleep loss in both male and female rats [63]. Importantly, pretreatment with a brain-penetrating inhibitor of kynurenine aminotransferase II (KAT II), the primary KYNA-synthesizing enzyme, prevents kynurenine-induced sleep disruption [62]. Consistent with this interpretation, mouse studies that disrupt KP metabolism by altering kynurenine 3-monooxygenase function and shift metabolism toward KYNA show altered sleep-wake architecture and further support a role for KYNA in sleep regulation [50]. Together, these findings support a causal role for KYNA in mediating sleep impairments following kynurenine exposure. Mechanistically, KYNA acts as an endogenous antagonist of α7 nicotinic acetylcholine receptors (α7nAChRs) and NMDA glutamate receptors [97]. Because REM sleep depends on high cholinergic tone [86], KYNA-mediated inhibition of α7nAChR signaling may contribute to the REM sleep deficits associated with elevated brain KYNA. KYNA also activates the aryl hydrocarbon receptor (AhR), which can influence circadian rhythms [98], and G-protein-coupled receptor 35 (GPR35), although the roles of these pathways in brain sleep regulation remain poorly understood, particularly during pregnancy.

Presently, we verified that a kynurenine diet increases brain KYNA levels irrespective of pregnancy status [22]. We observed elevated KYNA in the brainstem, cortex, hippocampus, and frontal cortex of both pregnant and non-pregnant females, indicating widespread neosynthesis from kynurenine. Because rodents predominantly consume food during the dark phase, measurements collected at ZT 6 likely provide a conservative estimate of diet-induced brain KYNA levels. While determining brain KYNA levels across a 24-h period and across days of diet administration could be informative, our current ZT 6 measurements confirm that KYNA remains elevated relative to matched controls during the primary sleep period in rodents, including during periods when REM sleep deficits were observed.

Regardless of kynurenine diet status, we observed elevated plasma kynurenine, KYNA, and QUIN levels in pregnant dams relative to non-pregnant females. These data corroborate existing literature showing that peripheral KP metabolism increases during gestation [32]. However, plasma KP measurements alone are not sufficient to determine the full contribution of downstream metabolites to maternal brain physiology. Although our mechanistic interpretation focuses on KYNA based on prior sleep studies and the observed elevations in brain KYNA, plasma QUIN was also increased during pregnancy and may indicate broader activation of KP metabolism. QUIN and other KP metabolites may contribute to aspects of maternal neurobiology or health beyond the REM sleep phenotype emphasized in our present findings.

In the pregnant brain, we observed elevated KYNA levels without a corresponding increase in brain kynurenine compared to non-pregnant females. Thus, increased kynurenine availability is unlikely to fully account for the greater brain KYNA accumulation observed during pregnancy. Instead, pregnancy may enhance kynurenine conversion to KYNA in brain astrocytes or alter the balance of KP metabolism among cell types. Future studies that measure KAT activity, kynurenine 3-monooxygenase activity, and cell-specific KP metabolism during pregnancy could further clarify these mechanisms. These studies will also help determine whether pregnancy-associated KP activation is an adaptive response that supports maternal physiology and fetal development, a consequence of fetal metabolic demands, or a vulnerability factor when excessively activated [27,32]. Nonetheless, KYNA levels were elevated in both pregnant and non-pregnant females to an extent associated with sleep disruption.

Importantly, we confirmed that neither pregnant nor non-pregnant kynurenine-fed females exhibited elevated corticosterone or broad peripheral inflammation, indicating that stress-related endocrine activation or systemic inflammation are unlikely to explain the observed sleep disturbances. Kynurenine diet did, however, modestly alter select circulating markers in non-pregnant females, increasing epidermal growth factor (EGF) while reducing interleukin-1 beta (IL-1β) and vascular endothelial growth factor (VEGF). These changes suggest that elevated KP metabolism may influence specific immune or growth-factor signaling pathways, although their relevance to KYNA-associated sleep disruption remains unclear.

### Postpartum sleep alterations after kynurenine exposure

Maternal sleep was also disrupted postpartum, although prior kynurenine exposure did not further alter gross sleep-wake architecture. In line with prior studies, rat dams showed reduced light phase NREM sleep and increased dark phase NREM and REM sleep postpartum [15,16]. Females were less able to maintain sleep episodes during the light phase, likely due in part to caregiving demands from pups [15]. However, increased dark phase NREM delta power indicates intact homeostatic sleep regulation and increased sleep drive during the dark phase, when total postpartum sleep increased relative to baseline. EKyn dams did not differ significantly from ECon dams in postpartum sleep-wake architecture, suggesting that the effects of kynurenine exposure on sleep duration and bout structure resolved after diet cessation and parturition. However, EKyn dams showed reduced postpartum NREM delta power and REM theta power, indicating that prior kynurenine exposure may have lingering effects on sleep quality that gross sleep-wake architecture alone does not capture.

### Implications for maternal and offspring health

Sleep during pregnancy is increasingly recognized as a critical determinant of maternal and offspring health. Poor maternal sleep is associated with postpartum depression, pregnancy complications, and altered neuropsychiatric development in offspring [3-10]. Changes to KP metabolism have also been implicated in many of these pregnancy complications and neurodevelopmental disorders [32-40]. Consistent with these clinical associations, rodent studies show that elevated prenatal KP metabolism, including in EKyn offspring, produces endophenotypes relevant to neurodevelopmental disorders [48-52,54-61]. Notably, EKyn offspring exhibit elevated brain KYNA, cognitive impairments, and sleep disturbances, features with relevance to schizophrenia and bipolar disorder [51-54,56-58]. Because prenatal insults associated with psychotic disorders also elevate KP metabolism [22,23,45,46,99-102], the EKyn model provides a valuable translational tool to study the effects of prenatal KP activation on offspring neurodevelopment. Our finding that the EKyn diet also disrupts maternal sleep raises the possibility that maternal sleep disturbance contributes to adverse maternal and offspring outcomes associated with perinatal KP activation.

Our study confirms that sleep changes dynamically across pregnancy and the postpartum period. We identified maternal sleep impairments in EKyn dams that were detectable only through within-subject comparisons, highlighting the importance of individualized assessment of sleep-wake phenotypes in identifying maternal sleep disturbances. Moreover, the immediate and persistent REM sleep deficits observed in non-pregnant females fed kynurenine indicate that pregnancy attenuates the sleep-disruptive effects of elevated KP metabolism. Together, these findings support prenatal KP dynamics as a modifiable neurobiological mechanism that may contribute to maternal sleep disturbances and adverse maternal and child health outcomes. Therapeutic approaches that improve maternal sleep, rebalance disrupted perinatal KP metabolism, or mitigate the effects of elevated KYNA may help promote long-term maternal and offspring health.

## Supporting information

Supplementary Statistical File

## Acknowledgments

The authors thank Snezana Milosavljevic, Sam C. Walther, and Corbin E. Witt for their technical contributions to this work. Mass spectrometry was performed at the University of South Carolina Mass Spectrometry Center [RRID:SCR_027213], supported by National Science Foundation grant 1828059. Some figure schematics were created with BioRender.com.

## Author Contributions

**CJW:** Conceptualization, investigation, methodology, formal analysis, visualization, writing – original draft, writing – review & editing, funding acquisition. **MVP:** methodology, writing – original draft, writing – review & editing. **JHC:** methodology, investigation, writing – review & editing. **HV:** software, writing – review & editing. **NF:** methodology, investigation, writing – review & editing, project administration, supervision, resources. **AP:** Conceptualization, investigation, methodology, formal analysis, visualization, writing – original draft, writing – review and editing, funding acquisition, project administration, supervision, resources.

