## Supplemental Figures for "Pregnancy Impacts Maternal Sleep: Focus on Kynurenine Pathway Activation in Female Rats"

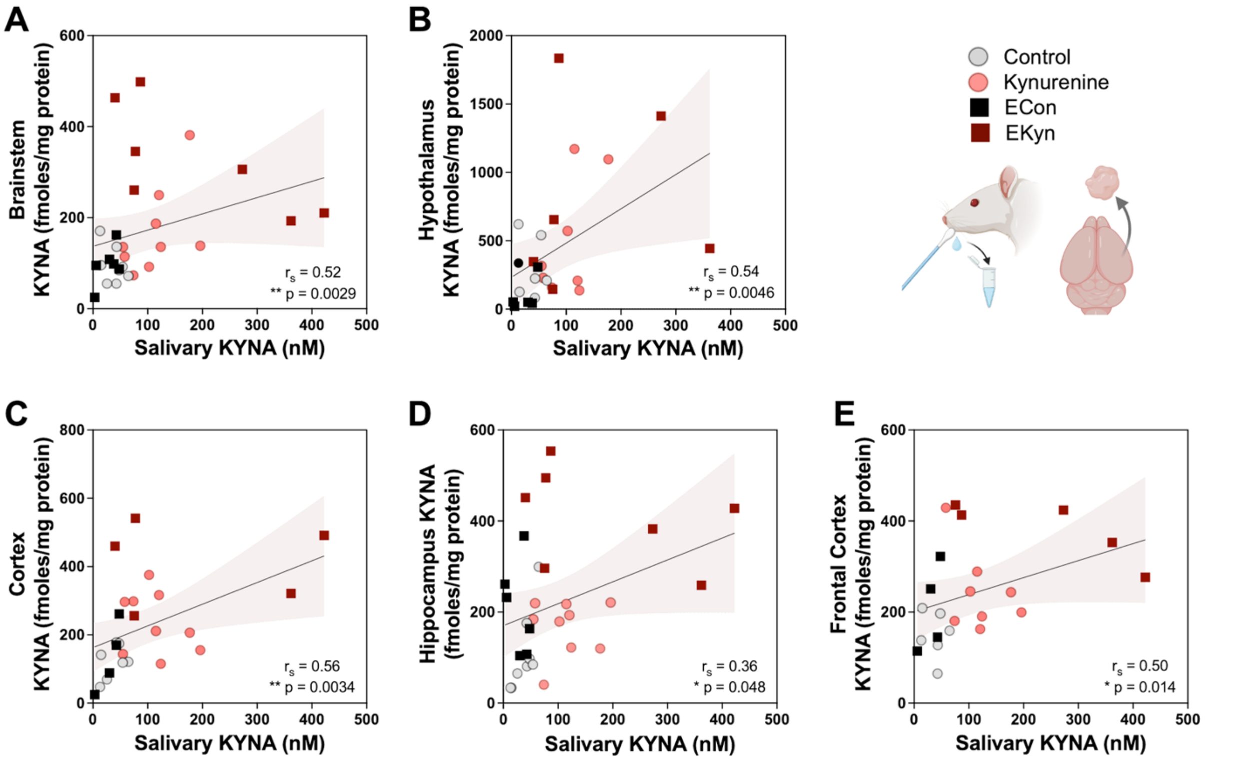


**Figure S1: Salivary KYNA positively correlates with brain KYNA content.** Pregnant and non-pregnant females received control diet or kynurenine-supplemented diet (+ 100 mg/day kynurenine). Correlations between salivary and **(A)** brainstem, **(B)** hypothalamus, **(C)** cortex, **(D)** hippocampus, and **(E)** frontal cortex KYNA. Spearman correlation: * $p < 0.05$, ** $p < 0.01$. N = 5-9/group.

**Kynurenine diet elevates KP metabolism without increasing plasma corticosterone or broadly altering inflammatory markers in non-pregnant females**

We confirmed that kynurenine diet in non-pregnant females elevates KP metabolism. As in previous studies with ECon and EKyn dams,[1] tissues were harvested at ZT 6 on day 7 of diet administration (**Figure S2A**). We first assessed plasma corticosterone to ensure kynurenine supplementation did not induce a stress response and found no effect of diet (**Figure S2B).** We also assessed a panel of inflammatory markers in plasma and observed only minor differences between groups (**Figure S3**). Specifically, kynurenine diet elevated plasma EGF ($p < 0.01$) yet reduced IL-1β ($p < 0.01$) and VEGF ($p < 0.05$). Mean plasma inflammatory marker concentrations and exact *p*-values are provided in Figure 2.

Kynurenine diet did not affect plasma tryptophan levels (**Figure S1C**), but increased plasma kynurenine 3.4-fold (**Figure S2D**, $p < 0.0001$), KYNA 2.3-fold (**Figure S2E**, $p < 0.0001$), and QUIN 2.2-fold (**Figure S2F**, $p < 0.05$). Kynurenine diet also elevated brain kynurenine levels compared with control across brain regions (**Figure S2G**: brainstem: +3.9-fold, $p < 0.01$; cortex: +2.7-fold, $p < 0.05$; hippocampus: +2-fold, $p < 0.05$; frontal cortex: +1.5-fold, $p < 0.05$) and KYNA levels compared to control (**Figure S2H**, brainstem: +1.9-fold; $p < 0.05$, cortex: +1.8-fold $p < 0.05$; hippocampus: +1.5-fold, $p < 0.05$; frontal cortex: + 1.6-fold, $p < 0.05$).

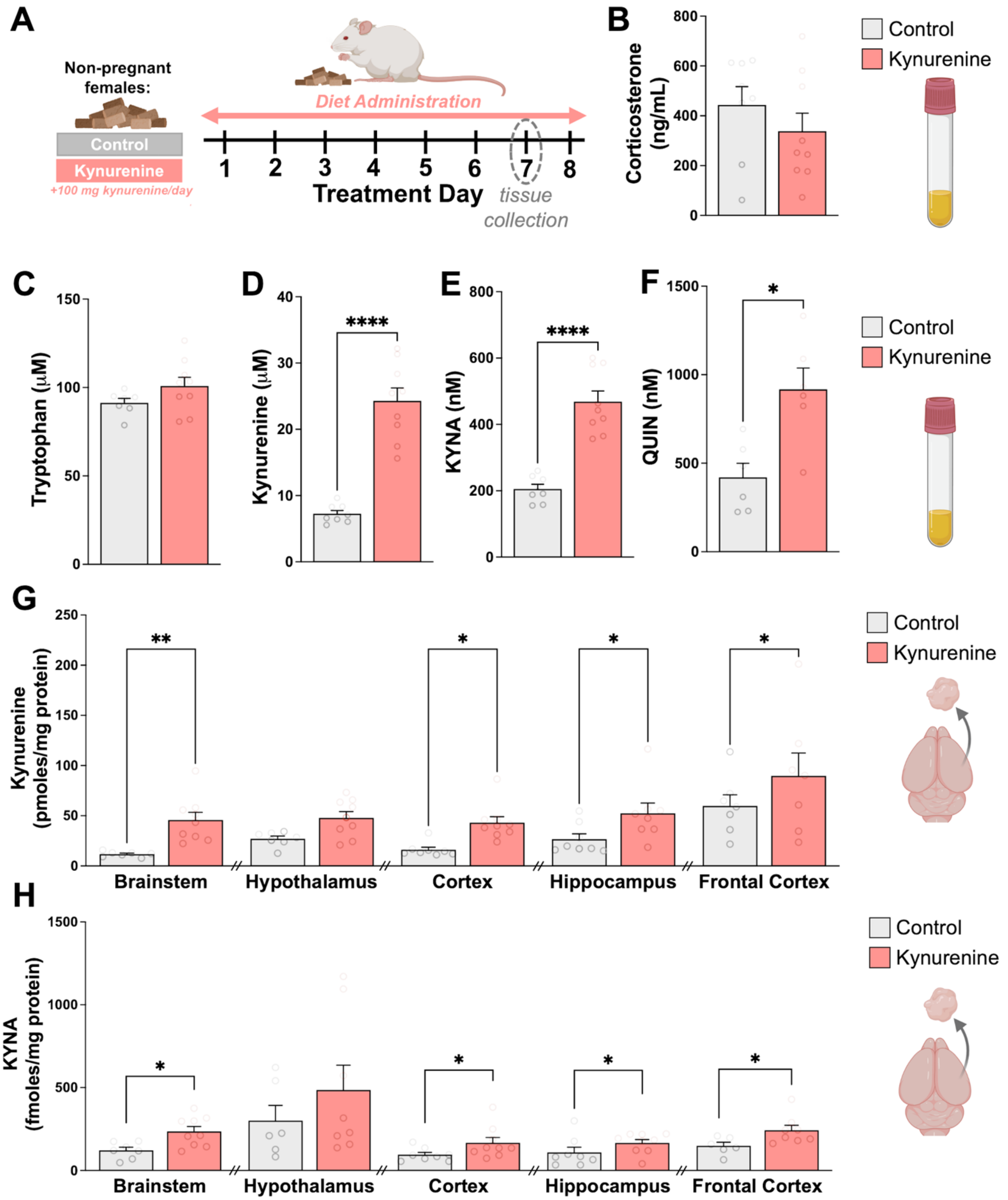


**Figure S2: Kynurenine diet elevates KP metabolites in plasma and brain of non-pregnant females. (A)** Experimental timeline. Non-pregnant females received daily control or kynurenine-supplemented diet (+ 100 mg/day kynurenine). Tissues were collected at ZT 6 on day 7 of diet administration. Plasma outcomes include **(B)** corticosterone, **(C)** tryptophan, **(D)** kynurenine, **(E)** KYNA, and **(F)** QUIN. Brain outcomes include **(G)** kynurenine and **(H)** KYNA. Data are mean ± SEM. Mann-Whitney tests: $p < 0.05$, ** $p < 0.01$, **** $p < 0.0001$. N=6-9/group.

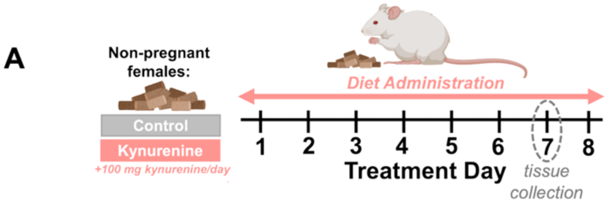


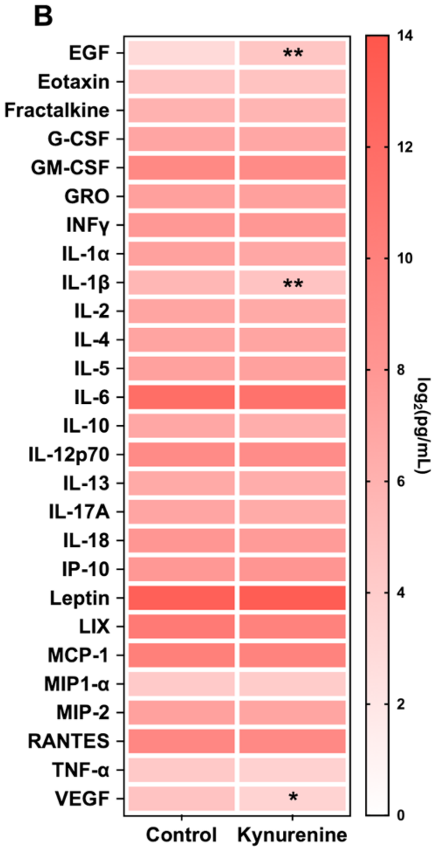


**C**

| Plasma Inflammatory Markers (pg/mL) | | | | |
|---|---|---|---|---|
| *Metabolite* | *Control (mean ± SEM)* | *Kynurenine (mean ± SEM)* | *Control vs Kynurenine Significance* | *p-value* |
| **EGF** | 8.9 ± 2.0 | 24.5 ± 6.4 | ** | 0.0043 |
| **Eotaxin** | 25.7 ± 1.5 | 26.5 ± 0.5 | ns | 0.5278 |
| **Fractalkine** | 63.0 ± 2.7 | 55.3 ± 2.5 | ns | 0.0657 |
| **G-CSF** | 125.6 ± 16.2 | 112.6 ± 7.2 | ns | 0.9003 |
| **GM-CSF** | 644.6 ± 69.3 | 591.3 ± 103.3 | ns | 0.8089 |
| **GRO** | 156.8 ± 9.7 | 157.4 ± 8.5 | ns | 0.9755 |
| **INF-γ** | 256.1 ± 50.8 | 242.3 ± 17.6 | ns | 0.8619 |
| **IL-1α** | 151.9 ± 14.8 | 116.8 ± 15.5 | ns | 0.0536 |
| **IL-1β** | 52.1 ± 5.27 | 27.1 ± 3.9 | ** | 0.0051 |
| **IL-2** | 134.3 ± 21.8 | 111.9 ± 21.0 | ns | 0.8112 |
| **IL-4** | 131.7 ± 11.71 | 128.8 ± 11.8 | ns | >0.9999 |
| **IL-5** | 153.0 ± 13.71 | 141.1 ± 8.2 | ns | 0.9268 |
| **IL-6** | 3805.0 ± 373.0 | 3653.0 ± 187.5 | ns | 0.7866 |
| **IL-10** | 122.2 ± 24.7 | 77.4 ± 6.4 | ns | 0.1389 |
| **IL-12p70** | 556.5 ± 43.6 | 465.3 ± 21.2 | ns | 0.1427 |
| **IL-13** | 98.9 ± 8.6 | 83.0 ± 10.0 | ns | 0.3129 |
| **IL-17A** | 122.7 ± 14.1 | 96.3 ± 13.0 | ns | 0.3462 |
| **IL-18** | 272.5 ± 43.8 | 209.2 ± 13.1 | ns | 0.3409 |
| **IP-10** | 240.4 ± 30.3 | 284.9 ± 58.9 | ns | 0.7308 |
| **leptin** | 8979.0 ± 1446.0 | 14182.0 ± 5628.0 | ns | 0.8357 |
| **LIX** | 2206.0 ± 744.7 | 991.4 ± 120.6 | ns | 0.2343 |
| **MCP-1** | 1179.0 ± 109.4 | 931.7 ± 95.0 | ns | 0.2343 |
| **MIP-1α** | 18.1 ± 1.5 | 16.7 ± 1.1 | ns | 0.5058 |
| **MIP-2** | 150.1 ± 15.5 | 129.0 ± 15.8 | ns | 0.5338 |
| **RANTES** | 774.6 ± 141.0 | 631.7 ± 90.7 | ns | 0.5338 |
| **TNF-α** | 19.0 ± 2.0 | 14.5 ± 2.4 | ns | 0.2768 |
| **VEGF** | 28.4 ± 5.7 | 12.7 ± 2.8 | * | 0.048 |

**Figure S3: Kynurenine diet minimally alters plasma inflammatory markers in non-pregnant females. (A)** Experimental timeline. Non-pregnant females received daily control or kynurenine-supplemented diet (+ 100 mg/day kynurenine). Tissues were collected at ZT 6 on day 7 of diet administration. **(B)** Plasma inflammatory markers measured using a 27-plex cytokine discovery panel. **(C)** Mean inflammatory marker concentrations ± SEM and Mann-Whitney test results: * $p$ < 0,05, ** $p$ < 0.01. N = 5-7/group.

**Pregnancy and kynurenine diet alter KP metabolites in female rats**

We next evaluated the effects of pregnancy on KP metabolites by comparing biochemical measurements from pregnant and non-pregnant females within control or kynurenine treatment conditions. KP metabolite levels were expressed as a percentage of non-pregnant control values. Pregnancy significantly affected plasma tryptophan levels ($F_{1, 28} = 37.14$, $p < 0.0001$) (**Figure S4B**), which were reduced by 32% in ECon dams ($p < 0.01$) and 40% in EKyn dams ($p < 0.0001$) compared to their non-pregnant counterparts. Main effects of pregnancy were also observed for plasma kynurenine ($F_{1, 27} = 4.919$, $p < 0.05$) (**Figure S4C**), KYNA ($F_{1, 27} = 12.21$, $p < 0.01$) (**Figure S4D**), and QUIN ($F_{1, 18} = 6.266$, $p < 0.05$) (**Figure S4E**). Post hoc analyses revealed significantly higher plasma kynurenine (+30%, $p < 0.05$) and KYNA (+53%, $p < 0.01$) in EKyn dams compared to non-pregnant kynurenine-treated females.

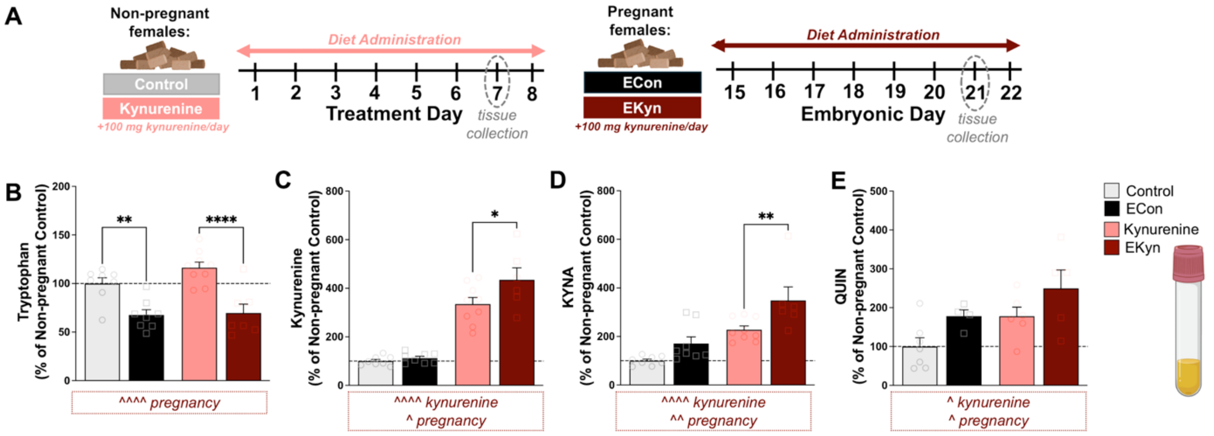


**Figure S4: Pregnancy influences basal and kynurenine-stimulated KP metabolites in female plasma. (A)** Experimental timeline. Pregnant and non-pregnant females received control diet or kynurenine-supplemented diet (+ 100 mg/day kynurenine). Tissues were collected at ZT 6 on day 7 of diet administration. Plasma outcomes include **(B)** tryptophan, **(C)** kynurenine, **(D)** KYNA, and **(E)** QUIN. Data are expressed as a percentage of non-pregnant control and shown as mean ± SEM. Two-way ANOVA. Significant main effects are displayed in boxed annotations below each figure panel: ^ $p < 0.05$, ^^ $p < 0.01$, ^^^^ $p < 0.0001$. Fisher's LSD post-hoc: * $p < 0.05$, ** $p < 0.01$, **** $p < 0.0001$ N = 4-9/group.

In the brain (**Figure S5**), pregnancy significantly affected kynurenine content only in the hypothalamus ($F_{1, 27}$ = 6.583, $p$ < 0.05) and cortex ($F_{1,30}$ = 4.817, $p$ < 0.05), where levels were increased by 98% and 43%, respectively, compared with non-pregnant counterparts (**Figure S5C, D**). Pregnancy also influenced KYNA content in the brainstem ($F_{1, 24}$ = 10.09, $p$ < 0.01) (**Figure S5G**), cortex ($F_{1, 27}$ = 7.973, $p$ < 0.01) (**Figure S5I**), hippocampus ($F_{1, 28}$ = 24.63, $p$ < 0.0001) (**Figure S5J**), and frontal cortex ($F_{1, 21}$ = 4.772, $p$ < 0.05) (**Figure S5K**). Compared with non-pregnant counterparts, ECon dams showed higher KYNA content in the brainstem (+140%, $p$ < 0.05) and hippocampus (+106%, $p$ < 0.05), whereas EKyn dams showed higher KYNA content in the brainstem (+76%, $p$ < 0.05), cortex (+94%, p < 0.01), hippocampus (+146%, $p$ < 0.0001), and frontal cortex (+99%, $p$ < 0.01).

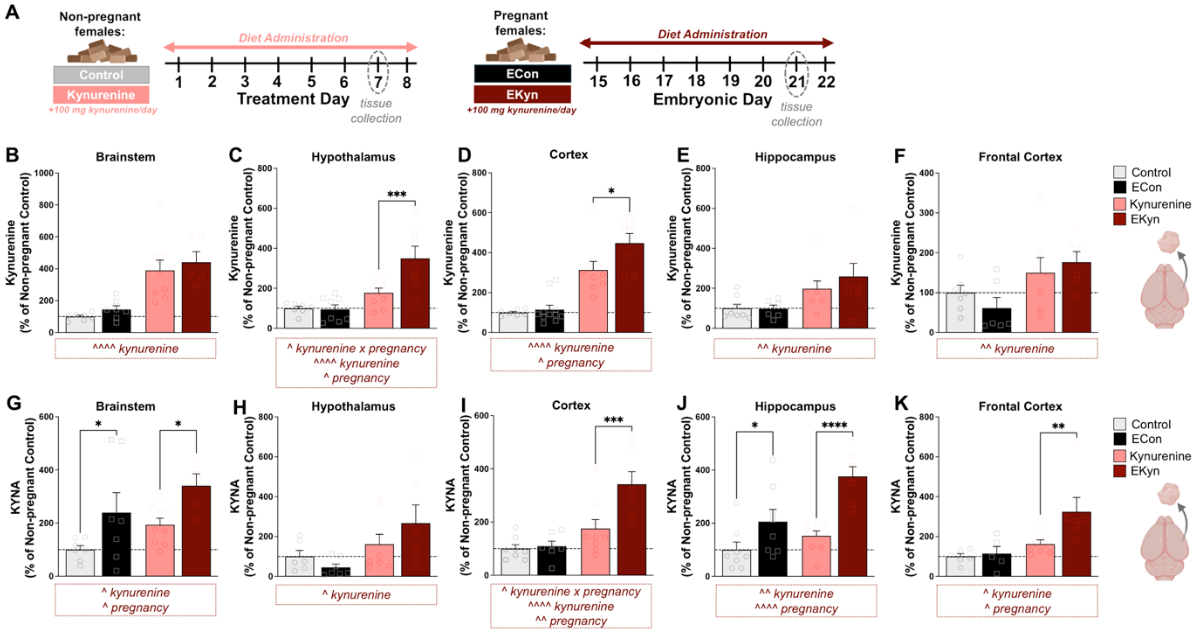


**Figure S5: Pregnancy influences basal and kynurenine-stimulated kynurenine and KYNA in the female brain. (A)** Experimental timeline. Pregnant and non-pregnant females received control diet or kynurenine-supplemented diet (+ 100 mg/day kynurenine). Tissues were collected at ZT 6 on day 7 of diet administration. Kynurenine was measured in the **(B)** brainstem, **(C)** hypothalamus, **(D)** cortex, **(E)** hippocampus, and **(F)** frontal cortex. KYNA was measured in the **(G)** brainstem, **(H)** hypothalamus, **(I)** cortex, **(J)** hippocampus, and **(K)** frontal cortex. Data are expressed as a percentage of non-pregnant control and shown as mean ± SEM. Two-way ANOVA. Significant main effects are displayed in boxed annotations below each figure panel: ^ $p$ < 0.05, ^^ $p$ < 0.01, ^^^^ $p$ < 0.0001. Fisher's LSD post hoc: * $p$ < 0.05, ** $p$ < 0.01, *** $p$ < 0.001, **** $p$ < 0.0001. N = 5-9/group.

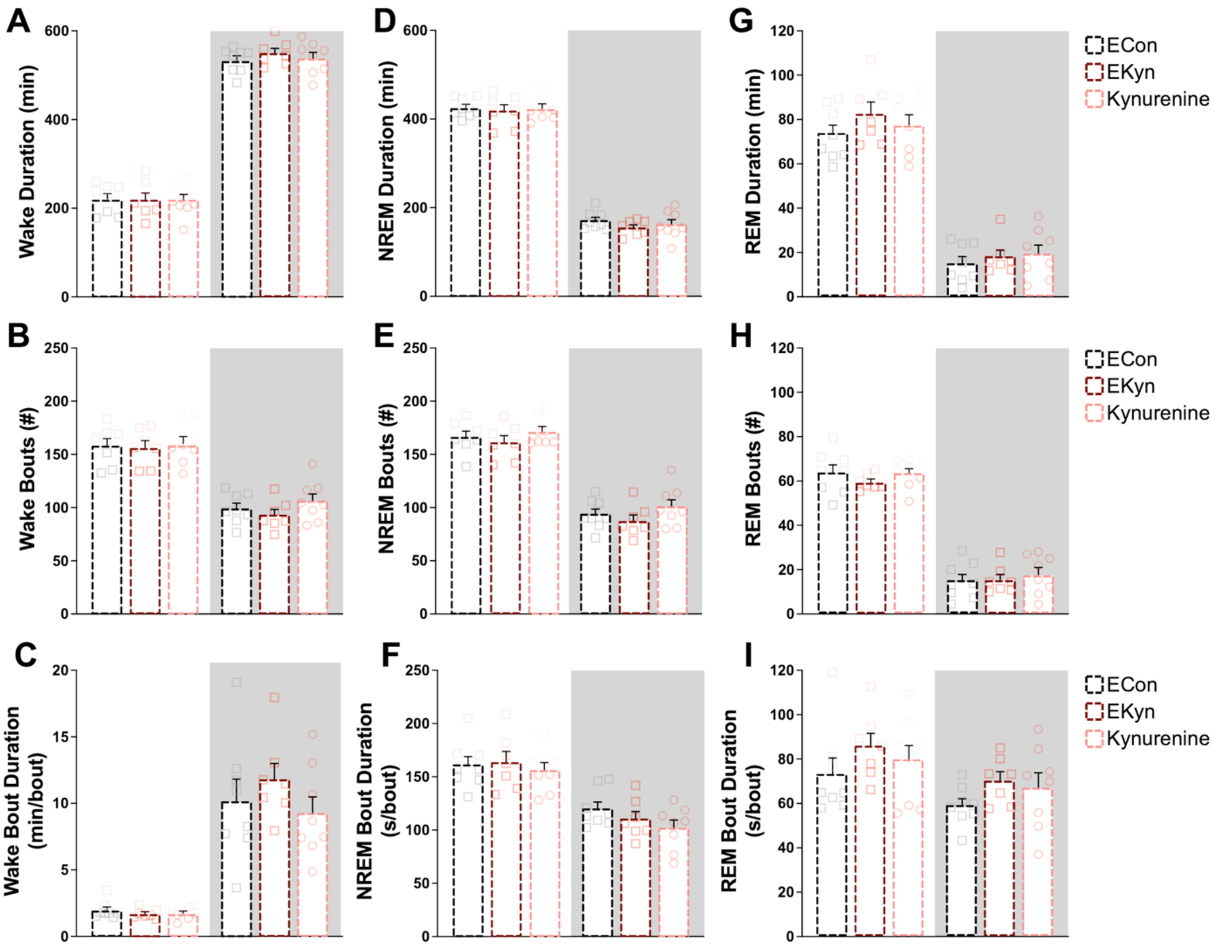


**Figure S6: Baseline sleep does not differ among ECon, EKyn, and kynurenine-treated cohorts.** Data are separated by light phase, shown with a white background, and dark phase, shown with a grey background. Wake outcomes include **(A)** duration, **(B)** bout number, and **(C)** bout duration. NREM sleep outcomes include **(D)** duration, **(E)** bout number, and **(F)** bout duration. REM sleep outcomes include **(G)** duration, **(H)** bout number, and **(I)** bout duration. Data are mean ± SEM. Two-way RM ANOVA. No statistical differences were detected. N = 7-8/group.

**Wakefulness, cage activity, and core body temperature are altered during pregnancy and postpartum**

During pregnancy, time spent awake was reduced during the dark phase (pregnancy x light phase interaction: $F_{2.988,\ 17.18} = 35.12$, $p < 0.0001$; main effect pregnancy: $F_{1,\ 7} = 619.4$, $p < 0.0001$) (**Figure S7A**). This reduction corresponded with the observed increases in NREM and REM sleep. On ED 20, ECon dams had more frequent wake episodes ($p < 0.05$) that were shorter in duration ($p < 0.05$) compared to baseline (**Figure S7B, C**). In line with reduced dark phase wakefulness, relative cage activity (main effect pregnancy: $F_{2.190,\ 15.33} = 3.99$, $p < 0.05$) and core body temperature (main effect pregnancy: $F_{2.344,\ 16.41} = 66.95$, $p < 0.0001$) were reduced across embryonic days in ECon dams (**Figure S7D, E**).

Postpartum wake timing was disrupted in ECon dams. Compared with baseline, wake duration was decreased during the light phase ($p < 0.01$) but increased during the dark phase ($p < 0.01$). Dark phase wake bouts were also more frequent ($p < 0.05$) and shorter in duration ($p < 0.05$), suggesting greater fragmentation of wakefulness during the active phase in the postpartum. Relative cage activity was not significantly altered postpartum; however, core body temperature was elevated during the light phase ($p < 0.0001$) and dark phase ($p < 0.01$). Effects of pregnancy and postpartum on sleep-wake transitions are presented in **Table S1**.

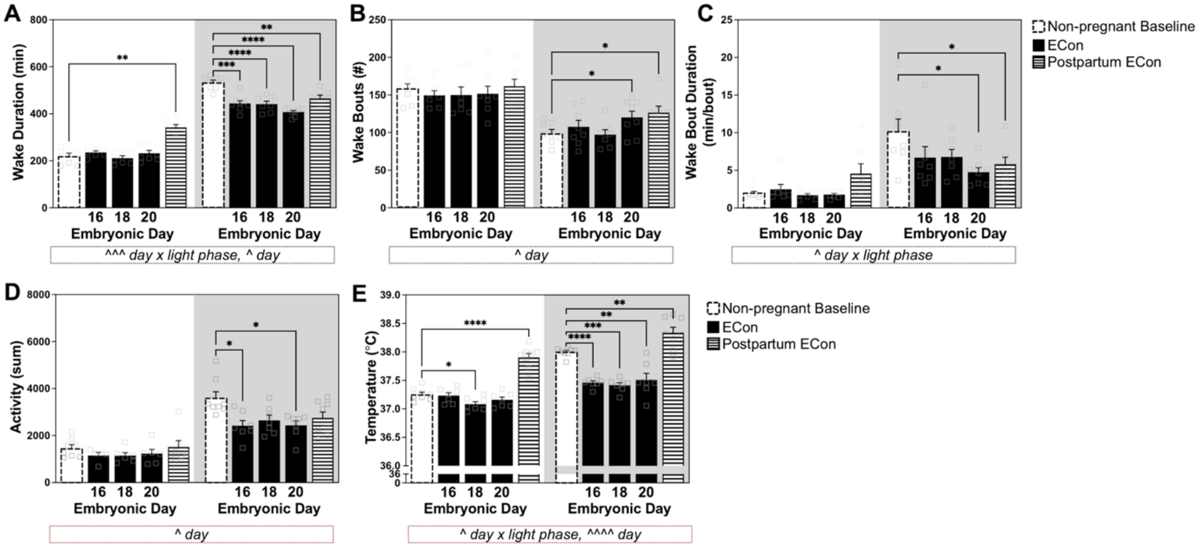


**Figure S7: Experimental day alters maternal wake architecture, cage activity, and body temperature in ECon dams**. Data are separated by light phase, shown with a white background, and dark phase, shown with a grey background. Wake outcomes include **(A)** duration, **(B)** bout number, and **(C)** bout duration. Additional outcomes include **(D)** relative cage activity and **(E)** core body temperature. Data are mean ± SEM. Two-way RM ANOVA. Significant interactions and main effects of day are displayed in boxed annotations below each figure panel: ^ $p < 0.05$, ^^^ $p < 0.001$, ^^^^ $p < 0.0001$. Fisher's LSD post-hoc: * $p < 0.05$, ** $p < 0.01$, *** $p < 0.001$, **** $p < 0.0001$. N = 5-8/group.

| Light Phase | | | | | |
|---|---|---|---|---|---|
| Treatment Day | Wake to NREM *Mean ± SEM* | NREM to Wake *Mean ± SEM* | NREM to REM *Mean ± SEM* | REM to Wake *Mean ± SEM* | REM to NREM *Mean ± SEM* |
| *Baseline* | 158.3 ± 6.1 | 98.2 ± 3.9 | 62.3 ± 3.9 | 59.7 ± 3.9 | 2.6 ± 0.6 |
| *ED 16* | 142.7 ± 7.3 | * 81.9 ± 6.5 | 64.6 ± 7.0 | 60.7 ± 6.5 | 3.7 ± 1.5 |
| *ED 18* | 146.8 ± 12.7 | * 82.3 ± 8.0 | 69.8 ± 7.8 | 64.3 ± 8.5 | 5.5 ± 1.2 |
| *ED 20* | 154.8 ± 11.3 | 86.2 ± 7.5 | 71.7 ± 7.3 | 68.5 ± 7.1 | 3.2 ± 0.5 |
| *PD 5* | 152.7 ± 10.2 | 109.5 ± 10.0 | * 47.8 ± 3.9 | ** 43.5 ± 4.5 | 4.3 ± 1.0 |
| **Dark Phase** | | | | | |
| **Treatment Day** | **Wake to NREM** *Mean ± SEM* | **NREM to Wake** *Mean ± SEM* | **NREM to REM** *Mean ± SEM* | **REM to Wake** *Mean ± SEM* | **REM to NREM** *Mean ± SEM* |
| *Baseline* | 91.1 ± 5.7 | 75.4 ± 5.1 | 15.9 ± 3.1 | 15.7 ± 3.0 | 0.18 ± 0.1 |
| *ED 16* | 95.1 ± 12.4 | 64.4 ± 7.9 | * 31.6 ± 7.8 | * 30.7 ± 7.5 | 1.0 ± 0.4 |
| *ED 18* | 92.2 ± 7.3 | * 63.3 ± 7.3 | ** 29.7 ± 3.6 | ** 29.0 ± 3.5 | 0.67 ± 0.4 |
| *ED 20* | * 115.2 ± 8.4 | 82.7 ± 8.4 | * 33.7 ± 6.0 | * 32.7 ± 6.2 | 1.0 ± 0.4 |
| *PD 5* | ** 125.3 ± 8.0 | 87.2 ± 4.6 | * 41.2 ± 10.5 | * 38.5 ± 10.4 | 2.7 ± 0.5 |

**Table S1: Pregnancy influences sleep-wake transitions.** Sleep-wake transitions were analyzed by two-way RM ANOVA. Fisher's LSD post hoc tests compared each pregnancy time point with baseline: * $p < 0.05$, ** $p < 0.01$. Data are mean ± SEM. N = 6-8/group/day.

**Wakefulness during pregnancy and postpartum is altered in EKyn dams**

Wake duration was reduced during the dark phase in EKyn dams, as indicated by a significant pregnancy x light phase interaction ($F_{1.972,\ 10.85} = 21.86$, $p < 0.001$) and main effect of pregnancy ($F_{2.941,\ 17.64} = 9.815$, $p < 0.001$) (**Figure S8A**). Significant pregnancy x light phase interactions also affected wake bout number ($F_{1.783,\ 9.808} = 4.797$, $p < 0.05$) (**Figure S8B**) and wake bout duration ($F_{2.108,\ 11.60} = 7.136$, $p < 0.01$) (**Figure S8C**). During the dark phase, wake bouts were increased on ED 16 ($p < 0.05$), whereas wake bout duration was decreased on ED 16 ($p < 0.05$) and ED 18 ($p < 0.05$). Relative cage activity (**Figure S8D**) and core body temperature (**Figure S8E**) were also reduced during the dark phase across embryonic days.

Postpartum, wakefulness increased during the light phase ($p < 0.01$) and decreased during the dark phase ($p < 0.0001$), mirroring postpartum changes in NREM and REM sleep. Despite reduced dark phase wake duration, wake bout numbers increased during the dark phase ($p < 0.01$), consistent with greater fragmentation of wakefulness. Core body temperature was increased during both the light phase ($p < 0.0001$) and dark phase ($p < 0.01$), whereas relative cage activity was reduced during the dark phase ($p < 0.01$). Effects of pregnancy and postpartum on EKyn sleep-wake transitions are presented in **Table S2**.

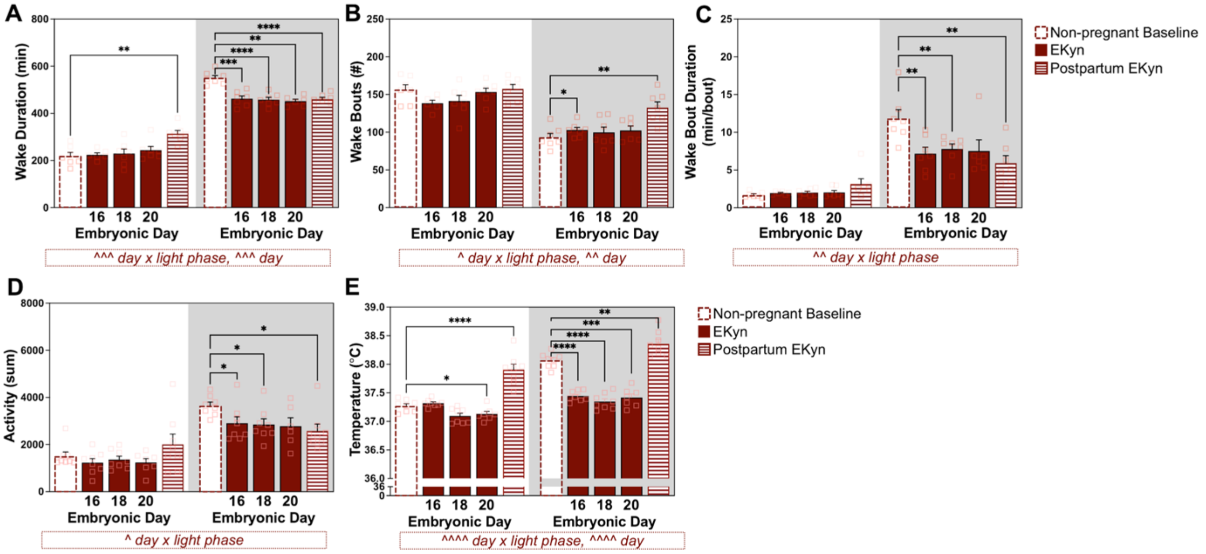


**Figure S8: Experimental day alters maternal wake architecture, cage activity, and body temperature in EKyn dams.** Data are separated by light phase, shown with a white background, and dark phase, shown with a grey background. Wake outcomes include **(A)** duration, **(B)** bout number, and **(C)** bout duration. Additional outcomes include **(D)** relative cage activity and **(E)** core body temperature. Data are mean ± SEM. Two-way RM ANOVA. Significant interactions and main effects of day are displayed in boxed annotation below each figure panel: ^ $p < 0.05$, ^^ $p < 0.01$, ^^^ $p < 0.001$, ^^^^ $p < 0.0001$. Fisher's LSD post-hoc: * $p < 0.05$, ** $p < 0.01$, *** $p < 0.001$, **** $p < 0.0001$. N = 7-8/group.

| Treatment Day | Wake to NREM | NREM to Wake | NREM to REM | REM to Wake | REM to NREM |
|---|---|---|---|---|---|
| **Light Phase** | | | | | |
| | *Mean ± SEM* | *Mean ± SEM* | *Mean ± SEM* | *Mean ± SEM* | *Mean ± SEM* |
| *Baseline* | 153.0 ± 6.4 | 96.6 ± 6.4 | 58.4 ± 1.2 | 56.1 ± 1.3 | 2.3 ± 0.5 |
| *ED 16* | * 135.7 ± 3.6 | * 77.4 ± 5.5 | 60.6 ± 3.6 | 58.1 ± 3.1 | 2.3 ± 0.7 |
| *ED 18* | 138.6 ± 7.7 | 85.3 ± 7.1 | 54.7 ± 2.6 | 52.9 ± 2.7 | 1.9 ± 0.7 |
| *ED 20* | 148.3 ± 4.8 | 96.9 ± 4.1 | 52.4 ± 2.6 | 51.1 ± 2.5 | * 1.1 ± 0.4 |
| *PD 5* | 153.3 ± 6.8 | ** 117.0 ± 4.0 | ** 40.3 ± 2.9 | *** 35.9 ± 2.8 | ** 4.4 ± 0.6 |
| **Dark Phase** | | | | | |
| **Treatment Day** | **Wake to NREM** | **NREM to Wake** | **NREM to REM** | **REM to Wake** | **REM to NREM** |
| | *Mean ± SEM* | *Mean ± SEM* | *Mean ± SEM* | *Mean ± SEM* | *Mean ± SEM* |
| *Baseline* | 84.3 ± 5.4 | 69.9 ± 3.8 | 15.1 ± 2.4 | 14.8 ± 2.3 | 0.32 ± 0.3 |
| *ED 16* | 87.3 ± 8.6 | 63.9 ± 4.8 | * 24.3 ± 4.8 | * 23.9 ± 4.6 | 0.57 ± 0.6 |
| *ED 18* | 91.3 ± 7.6 | 63.7 ± 6.0 | ** 28.9 ± 2.8 | ** 28.1 ± 2.7 | 0.71 ± 0.7 |
| *ED 20* | 94.7 ± 5.7 | 69.7 ± 6.0 | * 25.5 ± 3.8 | * 25.5 ± 3.6 | * 0.17 ± 0.2 |
| *PD 5* | * 124.4 ± 7.6 | * 96.1 ± 7.7 | **** 31.7 ± 2.5 | **** 28.7 ± 2.7 | ** 3.14 ± 1.0 |

**Table S2: EKyn influences sleep-wake transitions during pregnancy.** Sleep-wake transitions were analyzed by two-way RM ANOVA. Fisher's LSD post hoc tests compared each pregnancy time point with baseline: * $p < 0.05$, ** $p < 0.01$. Data are mean ± SEM. N = 7.

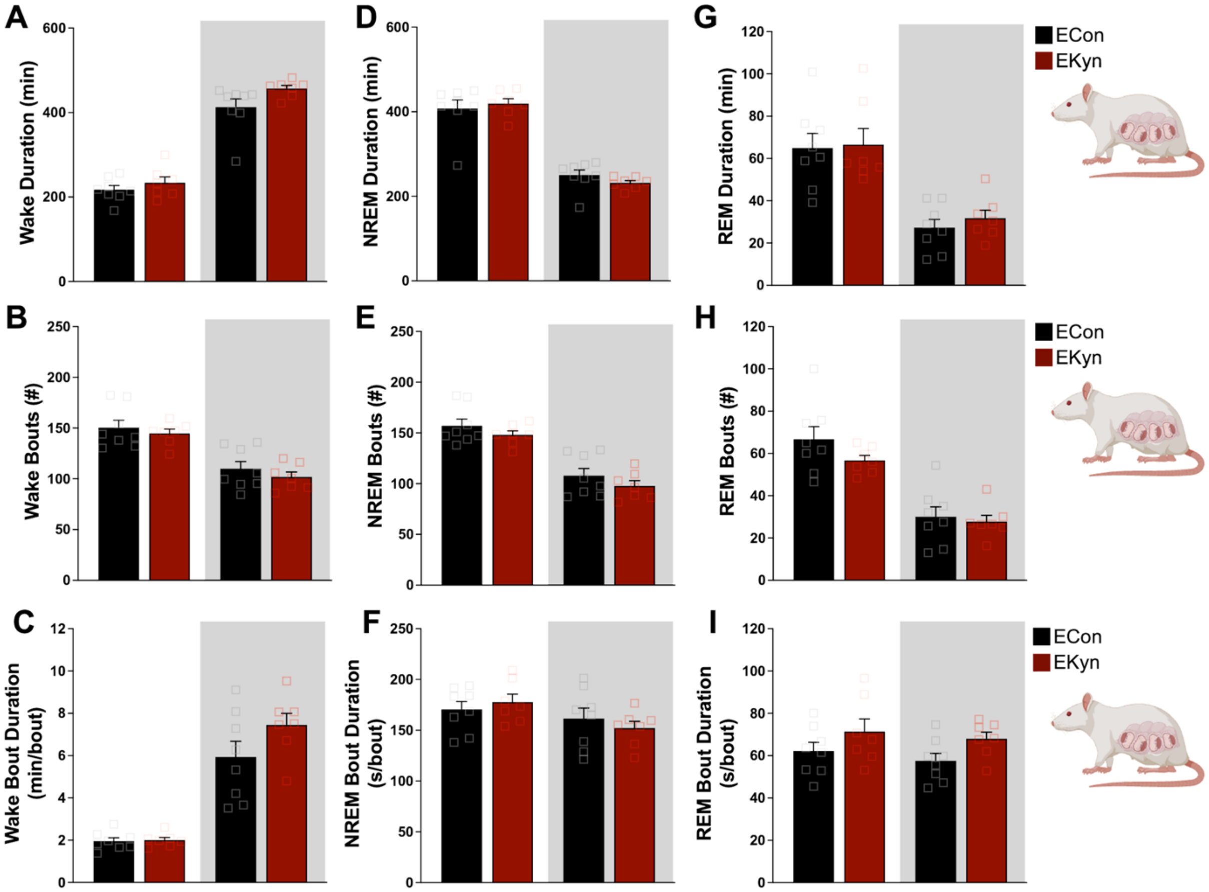


**Figure S9: Sleep-wake architecture does not differ between ECon and EKyn dams during the last week of gestation.** Data are separated by light phase, shown with a white background, and dark phase, shown with a grey background. Wake outcomes include **(A)** duration, **(B)** bout number, and **(C)** bout duration. NREM sleep outcomes include **(D)** duration, **(E)** bout number, and **(F)** bout duration. REM sleep outcomes include **(G)** duration, **(H)** bout number, and **(I)** bout duration. Data represent the average of treatment days and are shown as mean ± SEM. Two-way RM ANOVA. No statistical differences were detected. N = 7-8/group.

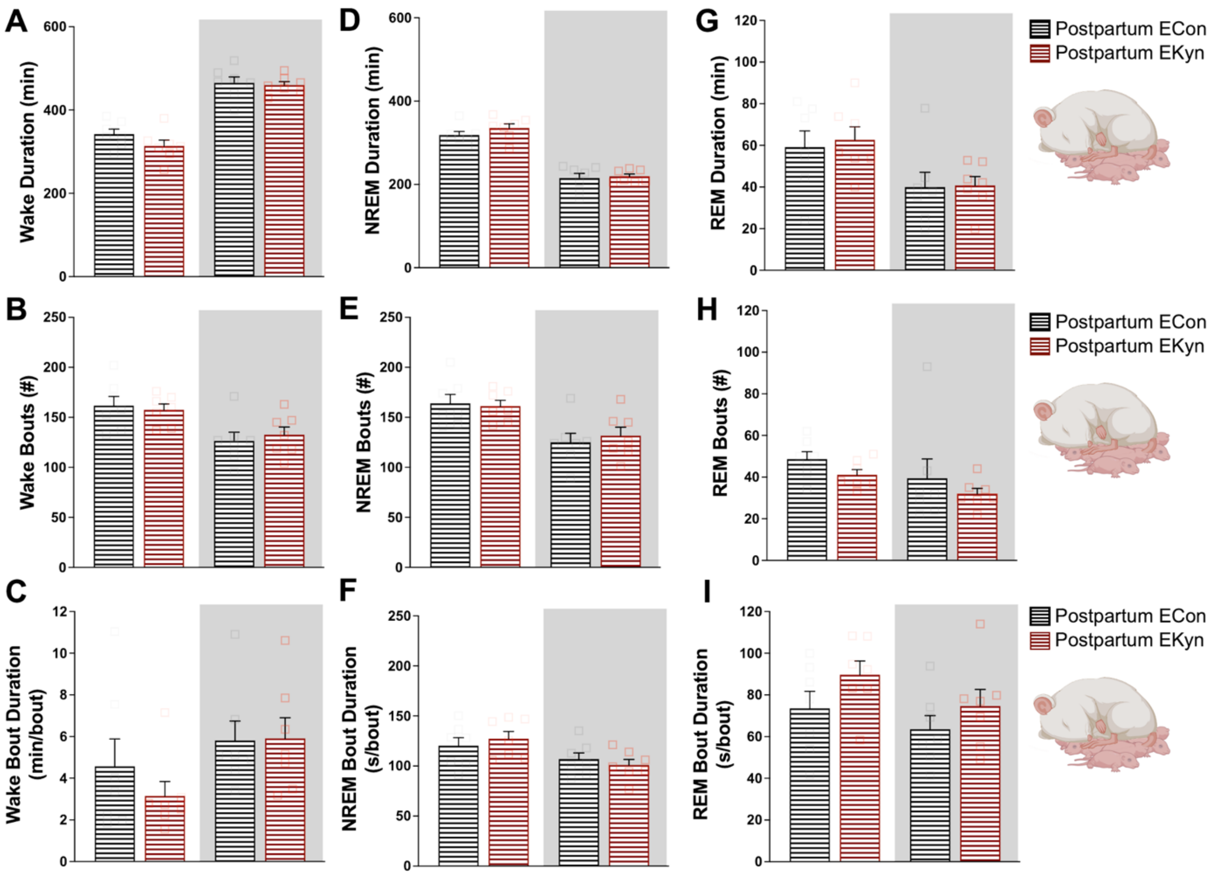


**Figure S10: Postpartum sleep-wake architecture does not differ between ECon and EKyn dams.** Data are separated by light phase, shown with a white background, and dark phase, shown with a grey background. Wake outcomes include **(A)** duration, **(B)** bout number, and **(C)** bout duration. NREM sleep outcomes include **(D)** duration, **(E)** bout number, and **(F)** bout duration. REM sleep outcomes include **(G)** duration, **(H)** bout number, and **(I)** bout duration. Data are mean ± SEM. Two-way RM ANOVA. No statistical differences were detected. N = 7/group.

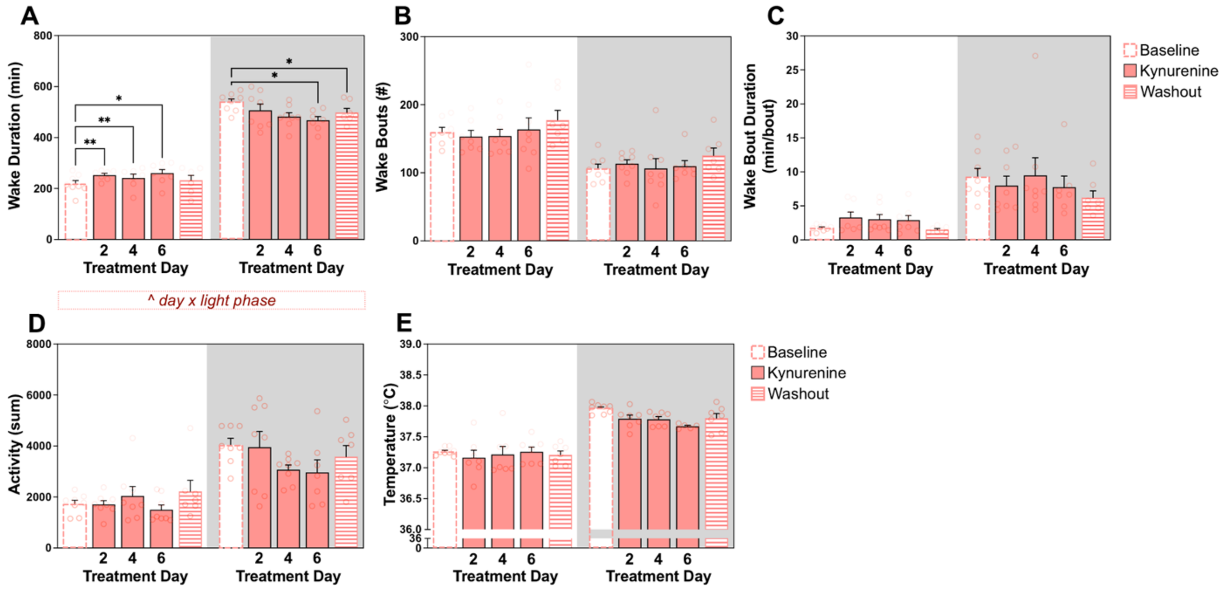


**Figure S11: Kynurenine diet does not affect NREM delta power or REM theta power in non-pregnant females.** Data are separated by light phase, shown with a white background, and dark phase, shown with a grey background. Spectral outcomes are shown during treatment **(A)** NREM sleep delta power, **(B)** REM sleep theta power, and washout **(C)** NREM sleep delta power, **(D)** REM sleep theta power compared to baseline. Data are mean ± SEM. Two-way RM ANOVA. No statistical differences were detected. N = 7-8/light phase.

**Kynurenine diet minimally alters wake architecture in non-pregnant females**

A significant kynurenine diet x light phase interaction affected wake duration ($F_{2.208,\ 12.7}$ = 5.432, $p < 0.05$) (**Figure S12A**). Time spent awake increased during the light phase on day 2 ($p < 0.01$), day 4 ($p < 0.01$), and day 6 ($p < 0.05$), but was reduced during the dark phase on day 6 ($p < 0.05$). Reduced dark phase wake duration persisted during washout ($p < 0.05$). Kynurenine diet did not significantly affect wake bout number (**Figure S12B**), wake bout duration (**Figure S12C**), relative cage activity (**Figure S12D**), or core body temperature (**Figure S12E**) in non-pregnant females.

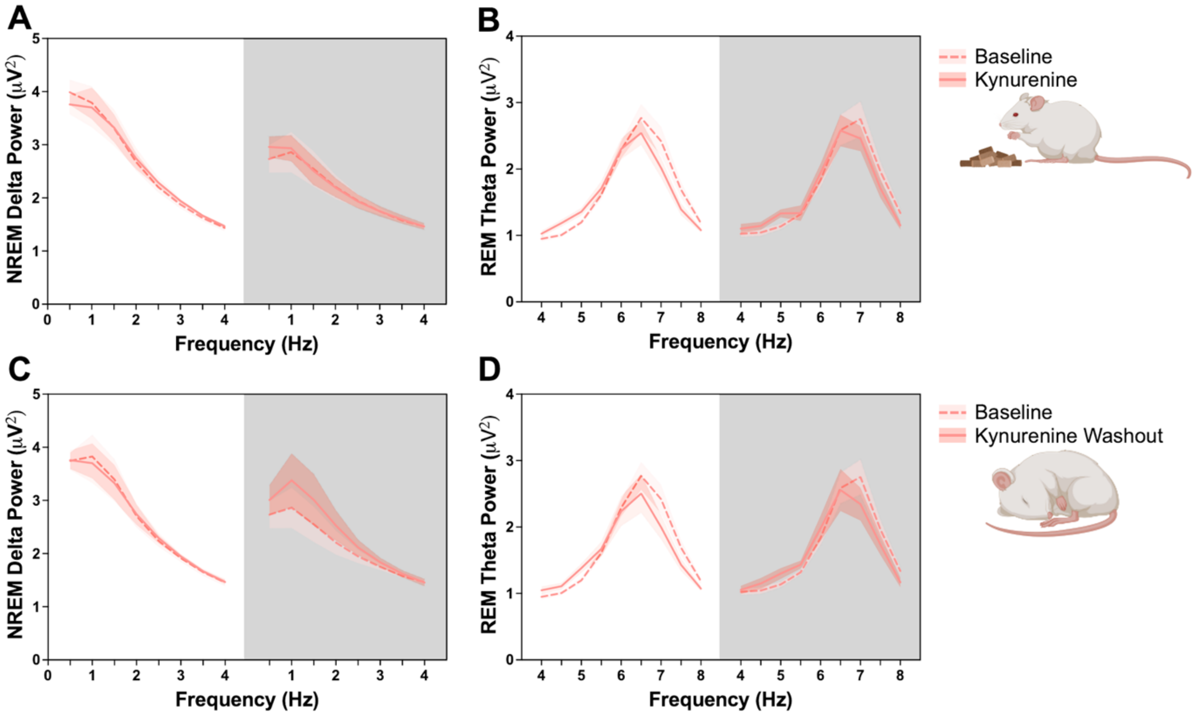


**Figure S12: Kynurenine diet minimally alters wake architecture without disrupting cage activity or body temperature in non-pregnant females.** Data are separated by light phase, shown in white background, and dark phase, shown in grey background. Wake outcomes include **(A)** duration, **(B)** bout number, and **(C)** bout duration. Additional outcomes include **(D)** relative cage activity and **(E)** core body temperature. Data are mean ± SEM. Two-way RM ANOVA. Significant interactions and main effects of treatment day are displayed in boxed annotation below each figure panel: ^ $p < 0.05$. Fisher's LSD post-hoc: * $p < 0.05$, ** $p < 0.01$. N = 7-8/day.

| Treatment Day | Light Phase<br>Wake to NREM<br>*Mean ± SEM* | NREM to Wake<br>*Mean ± SEM* | NREM to REM<br>*Mean ± SEM* | REM to Wake<br>*Mean ± SEM* | REM to NREM<br>*Mean ± SEM* |
|---|---|---|---|---|---|
| *Baseline* | 160.1 ± 6.9 | 102.8 ± 4.7 | 59.8 ± 4.1 | 59.0 ± 2.8 | 2.7 ± 0.7 |
| *Day 2* | 145.9 ± 8.4 | 91.9 ± 5.8 | 55.8 ± 4.2 | 54.0 ± 4.3 | 1.8 ± 0.6 |
| *Day 4* | 153.1 ± 8.6 | 108.5 ± 11.6 | 46.3 ± 4.3 | 44.6 ± 4.1 | 1.6 ± 0.6 |
| *Day 6* | 150.5 ± 8.1 | 96.8 ± 6.8 | 54.8 ± 4.2 | 53.8 ± 4.3 | 1.0 ± 0.3 |
| *Washout* | 172.3 ± 14.3 | 106.1 ± 9.4 | 67.3 ± 5.2 | 66.1 ± 5.3 | 1.0 ± 0.4 |
| **Treatment Day** | **Dark Phase**<br>**Wake to NREM**<br>*Mean ± SEM* | **NREM to Wake**<br>*Mean ± SEM* | **NREM to REM**<br>*Mean ± SEM* | **REM to Wake**<br>*Mean ± SEM* | **REM to NREM**<br>*Mean ± SEM* |
| *Baseline* | 96.5 ± 7.3 | 79.1 ± 7.1 | 18.2 ± 3.0 | 17.8 ± 2.9 | 0.63 ± 0.24 |
| *Day 2* | 105.4 ± 5.9 | 87.4 ± 8.2 | 18.4 ± 6.4 | 18.1 ± 6.2 | 0.25 ± 0.16 |
| *Day 4* | 104.9 ± 12.9 | 82.4 ± 8.9 | 23.0 ± 4.8 | 22.5 ± 4.9 | 0.50 ± 0.27 |
| *Day 6* | 112.1 ± 12.9 | 87.9 ± 12.5 | 24.8 ± 6.5 | 24.4 ± 6.4 | 0.38 ± 0.38 |
| *Washout* | 117.3 ± 11.4 | 93.3 ± 12.9 | 24.1 ± 5.3 | 24.0 ± 5.3 | 0.14 ± 0.14 |

**Table S3: Kynurenine diet does not influence sleep-wake transitions in non-pregnant females.** Data are mean ± SEM. Two-way RM ANOVA. No statistical differences were detected. N = 7-8/day.

| | | NREM Sleep Onset (Min) *Mean ± SEM* | | | | Two-way RM ANOVA |
|---|---|---|---|---|---|---|
| *Non-Pregnant Baseline* | *Treatment Group* | *Day 2* | *Day 4* | *Day 6* | *Washout* | *Results* |
| 9.4 ± 2.3 | *Kynurenine* | 16.9 ± 7.4 | 5.0 ± 2.0 | 3.9 ± 2.1 | 2.3 ± 1.1 | n.s. |
| | | *ED16* | *ED18* | *ED20* | *PD5* | |
| 9.6 ± 1.3 | *ECon* | 20.5 ± 4.9 | 5.3 ± 1.8 | 6.5 ± 2.9 | 7.8 ± 6.9 | n.s. |
| 10.2 ± 1.7 | *EKyn* | 14.1 ± 4.4 | 11.4 ± 5.1 | 5.9 ± 2.6 | 19.3 ± 7.3 | n.s. |
| | | **REM Sleep Onset (Min)** *Mean ± SEM* | | | | |
| *Non-Pregnant Baseline* | *Treatment Group* | *Day 2* | *Day 4* | *Day 6* | *Washout* | |
| 28.5 ± 4.3 | *Kynurenine* | 37.7 ± 10.9 | 18.1 ± 4.1 | 17.9 ± 3.6 | 12.81 ± 3 | n.s. |
| | | *ED16* | *ED18* | *ED20* | *PD5* | |
| 29.6 ± 2.2 | *ECon* | 44.9 ± 6.4 | 25.1 ± 4.7 | 37.2 ± 10.0 | 32.9 ± 12.5 | n.s. |
| 30.6 ± 3.7 | *EKyn* | 29.9 ± 7.3 | 36.6 ± 9.1 | 44.9 ± 18.2 | 65.9 ± 18.9 | n.s. |

**Table S4: NREM and REM sleep onset latencies are unaffected by treatment condition.** Data are mean ± SEM. Two-way RM ANOVA. No statistical differences were detected. N = 6-9/group.
