## Supplementary Statistical File for "Pregnancy Impacts Maternal Sleep: Focus on Kynurenine Pathway Activation in Female Rats"

| Figure | Statistical Test | Interaction | Results | Post-hoc Results (Fisher's LSD): |
|---|---|---|---|---|
| **1C** | 2-Way RM ANOVA | EKyn x day<br>EKyn<br>day | $F$ (1.107, 12.73) = 0.4125, $p$ = 0.5479<br>* $F$ (1, 12) = 7.301, $p$ = 0.0192<br>$F$ (1.107, 12.73) = 0.7786, $p$ = 0.4068 | *ECon vs EKyn:*<br>* ED 21: $p$ < 0.05 |
| **1D** | 2-Way RM ANOVA | kynurenine x day<br>kynurenine<br>day | $F$ (1.661, 23.25) = 1.369, $p$ = 0.2706<br>*** $F$ (1, 15) = 17.18, $p$ = 0.0009<br>$F$ (1.661, 23.25) = 2.131, $p$ = 0.1477 | *Control vs Kynurenine:*<br>* day 1: $p$ < 0.05<br>** day 4: $p$ < 0.01<br>* day 6: $p$ < 0.05 |
| **2B** | 2-Way RM ANOVA | pregnancy x light phase<br>pregnancy<br>light phase | **** $F$ (1.809, 10.85) = 30.72, $p$ < 0.0001<br>****$F$ (2.547, 17.83) = 19, $p$ < 0.0001<br>**** $F$ (1, 7) = 791,7, $p$ < 0.0001 | *Baseline vs ECon:*<br>* ED 18 (light phase): $p$ < 0.05<br>*** ED 16 (dark phase): $p$ < 0.001<br>**** ED 18 (dark phase): $p$ < 0.0001<br>**** ED 20 (dark phase): $p$ < 0.0001 |
| **2C** | 2-Way RM ANOVA | pregnancy x light phase<br>pregnancy<br>light phase | * $F$ (1.948, 11.69) = 5.663, $p$ = 0.0196<br>$F$ (2.173, 15.21) = 3.018, $p$ = 0.0754<br>**** $F$ (1, 7) = 269.2, $p$ < 0.0001 | *Baseline vs ECon:*<br>* ED 16 (light phase): $p$ < 0.05<br>** ED 20 (dark phase): $p$ < 0.01) |
| **2D** | 2-Way RM ANOVA | pregnancy x light phase<br>pregnancy<br>light phase | $F$ (2.405, 14.43) = 3.267, $p$ = 0.0604<br>** $F$ (1.664, 11.65) = 8.026, $p$ = 0.0083<br>* $F$ (1, 7) = 8.311, $p$ = 0.0236 | *Baseline vs ECon:*<br>** ED 16 (dark phase): $p$ < 0.01<br>** ED 18 (dark phase): $p$ < 0.01 |
| **2E** | 2-Way RM ANOVA | light phase:<br>pregnancy x frequency<br>pregnancy<br>frequency | * $F$ (1.0, 4.857) = 7.778, $p$ = 0.0397<br>$F$ (1, 6) = 0.0683, $p$ = 0.8025<br>**** $F$ (1.897, 11.38) = 80.85, $p$ < 0.0001 | *Baseline vs ECon:*<br>** 0.5 Hz: $p$ < 0.01<br>* 3 Hz: $p$ < 0.05<br>* 3.5 Hz: $p$ < 0.05<br>* 4 Hz: $p$ < 0.05 |
| | | dark phase:<br>pregnancy x frequency<br>pregnancy<br>frequency | $F$ (0.9565, 4.646) = 3.713, $p$ = 0.1166<br>* $F$ (1, 6) = 9.916, $p$ = 0.0198<br>**** $F$ (1.791, 10.75) = 37.81, $p$ < 0.0001 | *Baseline vs ECon:*<br>* 1 Hz: $p$ < 0.05<br>* 1.5 Hz: $p$ < 0.01<br>* 2.5 Hz: $p$ < 0.05<br>* 3 Hz: $p$ < 0.05 |
| **2F** | 2-Way RM ANOVA | pregnancy x light phase<br>pregnancy<br>light phase | * $F$ (1.789, 10.73) = 6.548, $p$ = 0.0156<br>$F$ (2.572, 18) = 0.9945, $p$ = 0.4074<br>**** $F$ (1, 7) = 159.2, $p$ < 0.0001 | *Baseline vs ECon:*<br>** ED 16 (dark phase): $p$ < 0.01<br>* ED 18 (dark phase): $p$ < 0.05<br>** ED 20 (dark phase): $p$ < 0.01 |
| **2G** | 2-Way RM ANOVA | pregnancy x light phase<br>pregnancy<br>light phase | $F$ (1.659, 9,955) = 2.538, $p$ = 0.1336<br>* $F$ (1.767, 19.37) = 5.065, $p$ = 0.0106<br>**** $F$ (1, 7) = 210.5, $p$ < 0.0001 | *Baseline vs ECon:*<br>** ED 16 (dark phase): $p$ < 0.01<br>** ED 18 (dark phase): $p$ < 0.01<br>** ED 20 (dark phase): $p$ < 0.01 |
| **2H** | 2-Way RM ANOVA | pregnancy x light phase<br>pregnancy<br>light phase | $F$ (2.217, 13.3) = 2.028, $p$ = 0.1677<br>$F$ (2.041, 14.28) = 1.522, $p$ = 0.2517<br>$F$ (1, 7) = 4.263, $p$ = 0.0778 | |
| **2I** | 2-Way RM ANOVA | light phase:<br>pregnancy x frequency<br>pregnancy<br>frequency | $F$ (0.1695, 0.8264) = 3.334, $p$ = 0.1794<br>* $F$ (1. 6) = 9.596, $p$ = 0.0212<br>* $F$ (1.327, 7.963) = 8.835, $p$ = 0.014 | *Baseline vs ECon:*<br>* 7 Hz: $p$ < 0.05<br>* 7.5 Hz: $p$ < 0.05 |
| | | dark phase:<br>pregnancy x frequency<br>pregnancy<br>frequency | $F$ (0.986, 4.807) = 1.392, $p$ = 0.2922<br>$F$ (1, 6) = 2.078, $p$ = 0.1996<br>* $F$ (1.22, 7.320) = 7.917, $p$ = 0.0216 | |
| **3D** | 2-Way RM ANOVA | light phase:<br>postpartum x frequency<br>postpartum<br>frequency | $F$ (1.441, 6.999) = 3.292, $p$ = 0.1059<br>$F$ (1, 6) = 3.457, $p$ = 0.1123<br>**** $F$ (1.817, 10.90) = 75.85, $p$ < 0.0001 | |
| | | dark phase:<br>postpartum x frequency<br>postpartum<br>frequency | * $F$ (0.9310, 4.522) = 7.398, $p$ = 0.0474<br>** $F$ (1, 6) = 25.33, $p$ = 0.0024<br>**** $F$ (1.845, 11.07) = 41.19, $p$ < 0.0001 | *Baseline vs postpartum ECon:*<br>* 1 Hz: $p$ < 0.05<br>*** 1.5 Hz: $p$ < 0.001<br>** 2 Hz: $p$ < 0.01<br>* 2.5 Hz: $p$ < 0.05 |

| Figure | Statistical Test | Interaction | Results | Post-hoc Results (Fisher's LSD): |
|---|---|---|---|---|
| **3E** | 2-Way RM ANOVA | light phase:<br>postpartum x frequency<br>postpartum<br>frequency | $F$ (0.3781, 1.843) = 6.976, $p$ = 0.1075<br>* $F$ (1, 6) = 12.60, $p$ = 0.0121<br>* $F$ (1.206, 7.235) = 8.844, $p$ = 0.0173 | *Baseline vs postpartum ECon:*<br>5 Hz: $p$ < 0.01<br>** 5.5 Hz: $p$ < 0.01<br>* 6 Hz: $p$ < 0.05<br>* 6.5 Hz: $p$ < 0.05 |
| | | dark phase:<br>postpartum x frequency<br>postpartum<br>frequency | $F$ (1.105, 5.389) = 0.6451, $p$ = 0.4715<br>$F$ (1, 6) = 4.381, $p$ = 0.0812<br>* $F$ (1.219, 7.317) = 8.995, $p$ = 0.0162 | |
| **4B** | 2-Way RM ANOVA | pregnancy x light phase<br>pregnancy<br>light phase | ** $F$ (1.628, 8.683) = 10.95, $p$ = 0.0054<br>** $F$ (2.107, 12.64) = 11.54, $p$ = 0.0013<br>**** $F$ (1, 6) = 366.9, $p$ < 0.0001 | *Baseline vs EKyn:*<br>**** ED 16 (dark phase): $p$ < 0.0001<br>**** ED 18 (dark phase): $p$ < 0.0001<br>**** ED 20 (dark phase): $p$ < 0.0001 |
| **4C** | 2-Way RM ANOVA | pregnancy x light phase<br>pregnancy<br>light phase | * $F$ (1.737, 9.266) = 7.148, $p$ = 0.0152<br>$F$ (2.083, 12.5) = 0.6213, $p$ = 0.5592<br>**** $F$ (1, 6) = 227.6, $p$ < 0.0001 | *Baseline vs EKyn:*<br>* ED 16 (light phase): $p$ < 0.05<br>* ED 16 (dark phase): $p$ < 0.05 |
| **4D** | 2-Way RM ANOVA | pregnancy x light phase<br>pregnancy<br>light phase | ** $F$ (1.830, 9.761) = 11.63, $p$ = 0.003<br>* $F$ (2.125, 12.75) = 3.936, $p$ = 0.0447<br>** $F$ (1, 6) = 18.06, $p$ = 0.0054 | *Baseline vs EKyn:*<br>* ED 16 (dark phase): $p$ < 0.05<br>** ED 18 (dark phase): $p$ < 0.01<br>** ED 20 (dark phase): $p$ < 0.01 |
| **4E** | 2-Way RM ANOVA | light phase:<br>pregnancy x frequency<br>pregnancy<br>frequency | $F$ (1.608, 9.647) = 0.1046, $p$ = 0.8619<br>$F$ (1, 6) = 0.2241, $p$ = 0.6527<br>**** $F$ (1.723, 10.34) = 49.58, $p$ < 0.0001 | |
| | | dark phase:<br>pregnancy x frequency<br>pregnancy<br>frequency | * $F$ (1.843, 8.949) = 6.609, $p$ = 0.0186<br>** $F$ (1, 6) = 28.15, $p$ = 0.0018<br>**** $F$ (1,797, 10.78) = 33.57, $p$ < 0.0001 | *Baseline vs EKyn:*<br>** 1 Hz: $p$ < 0.01<br>** 1.5 Hz: $p$ < 0.01<br>*** 2 Hz: $p$ < 0.001<br>* 2.5 Hz: $p$ < 0.05<br>** 3 Hz: $p$ < 0.001<br>* 3.5 Hz: $p$ < 0.05 |
| **4F** | 2-Way RM ANOVA | pregnancy x light phase<br>pregnancy<br>light phase | *** $F$ (1.871, 9.978) = 16.32, $p$ = 0.0008<br>$F$ (1.773, 10.64) = 2.201, $p$ = 0.1611<br>*** $F$ (1, 6) = 50.29, $p$ = 0.0004 | *Baseline vs EKyn:*<br>* ED 16 (light phase): $p$ < 0.05<br>** ED 18 (light phase): $p$ < 0.01<br>** ED 16 (dark phase): $p$ < 0.01<br>** ED 18 (dark phase): $p$ < 0.01<br>* ED 20 (dark phase): $p$ < 0.05 |
| **4G** | 2-Way RM ANOVA | pregnancy x light phase<br>pregnancy<br>light phase | ** $F$ (1.744, 0.303) = 8.574, $p$ = 0.009<br>* $F$ (1.674, 10.05) = 5.013, $p$ = 0.0353<br>**** $F$ (1, 6) = 112.6, $p$ < 0.0001 | *Baseline vs EKyn:*<br>*** ED 16 (dark phase): $p$ < 0.001<br>** ED 18 (dark phase): $p$ < 0.01<br>* ED 20 (dark phase): $p$ < 0.05 |
| **4H** | 2-Way RM ANOVA | pregnancy x light phase<br>pregnancy<br>light phase | $F$ (1.922, 10.25) = 3.418, $p$ = 0.0742<br>$F$ (1.762, 10.57) = 2.107, $p$ = 0.1721<br>$F$ (1, 6) = 1.281, $p$ = 0.3009 | |
| **4I** | 2-Way RM ANOVA | light phase:<br>pregnancy x frequency<br>pregnancy<br>frequency | $F$ (1.425, 8.552) = 3.838, $p$ = 0.0743<br>$F$ (1, 6) = 0.5181, $p$ = 0.4987<br>** $F$ (1.231, 7.385) = 15.00, $p$ = 0.0037 | |
| | | dark phase:<br>pregnancy x frequency<br>pregnancy<br>frequency | * $F$ (1.658, 8.082) = 4.758, $p$ = 0.047<br>$F$ (1, 6) = 0.6852, $p$ = 0.4395<br>** $F$ (1.222, 7.332) = 15.21, $p$ = 0.0043 | *Baseline vs EKyn:*<br>* 5.5 Hz: $p$ < 0.05<br>* 7.5 Hz: $p$ < 0.05<br>** 8 Hz: $p$ < 0.01 |
| **5D** | 2-Way RM ANOVA | light phase:<br>postpartum x frequency<br>postpartum<br>frequency | * $F$ (1.671, 10.03) =4.554, $p$ = 0.0441<br>* $F$ (1, 6) = 7.960, $p$ = 0.0303<br>**** $F$ (1.689, 10.14) = 51.15, $p$ < 0.0001 | *Baseline vs postpartum EKyn:*<br>* 1.5 Hz: $p$ < 0.05<br>** 2 Hz: $p$ < 0.01 |
| | | dark phase:<br>postpartum x frequency<br>postpartum<br>frequency | $F$ (2.207, 13.24) = 3.541, $p$ = 0.0551<br>* $F$ (1, 6) = 6.167, $p$ = 0.0476<br>**** $F$ (2.148, 12.89) = 40.26, $p$ < 0.0001 | *Baseline vs postpartum EKyn:*<br>* 1 Hz: $p$ < 0.05<br>* 1.5 Hz: $p$ < 0.05 |

| Figure | Statistical Test | Interaction | Results | Post-hoc Results (Fisher's LSD): |
|---|---|---|---|---|
| **5E** | 2-Way RM ANOVA | light phase:<br>postpartum x frequency<br>postpartum<br>frequency | * $F$ (1.874, 11.25) = 5.409, $p$ = 0.0240<br>* $F$ (1, 6) = 7.684, $p$ = 0.0323<br>** $F$ (1.162, 6.974) = 16.22, $p$ = 0.0043 | *<u>Baseline vs postpartum EKyn:</u>*<br>** 5.5 Hz: $p$ < 0.01<br>** 6 Hz: $p$ < 0.01 |
| | | dark phase:<br>postpartum x frequency<br>postpartum<br>frequency | $F$ (1.644, 9.862) = 2.861, $p$ = 0.1107<br>* $F$ (1, 6) = 7.277, $p$ = 0.0357<br>** $F$ (1.160, 6.961) = 20.06, $p$ = 0.0024 | *<u>Baseline vs postpartum EKyn:</u>*<br>* 7 Hz: $p$ < 0.05<br>* 7.5 Hz: $p$ < 0.05<br>* 8 Hz: $p$ < 0.05 |
| **6B** | 2-Way RM ANOVA | day x light phase<br>day<br>light phase | * $F$ (2.462, 14.16) = 4.457, $p$ = 0.0259<br>$F$ (2.320, 16.24) = 1.353, $p$ = 0.2891<br>**** $F$ (1, 7) = 362.4, $p$ < 0.0001 | *<u>Baseline vs Kynurenine:</u>*<br>day 4 (dark phase): $p$ < 0.05<br>day 6 (dark phase): $p$ < 0.01<br>washout (dark phase): $p$ < 0.05 |
| **6C** | 2-Way RM ANOVA | day x light phase<br>day<br>light phase | $F$ (1,588, 9.526) = 1.482, $p$ = 0.2695<br>$F$ (2.855, 19.99) = 1.690, $p$ = 0.2026<br>**** $F$ (1, 7) = 142, $p$ < 0.0001 | |
| **6D** | 2-Way RM ANOVA | day x light phase<br>day<br>light phase | $F$ (2.314, 13.88) = 2.635, $p$ = 0.1015<br>$F$ (2.883, 20.18) = 1.820, $p$ = 0.1771<br>*** $F$ (1, 7) = 40.74, $p$ = 0.0004 | |
| **6E** | 2-Way RM ANOVA | day x light phase<br>day<br>light phase | * $F$ (1.377, 8.264) = 5.844, $p$ = 0.0339<br>$F$ (2.513, 17.59) = 1.833, $p$ = 0.1836<br>** $F$ (1, 7) = 38, $p$ = 0.0005 | *<u>Baseline vs Kynurenine:</u>*<br>* day 2 (light phase): $p$ < 0.05<br>*** day 4 (light phase): $p$ < 0.001<br>* day 6 (light phase): $p$ < 0.05 |
| **6F** | 2-Way RM ANOVA | day x light phase<br>day<br>light phase | $F$ (1.327, 7.962) = 2.676, $p$ = 0.1374<br>$F$ (2.469, 17.28) = 1.599, $p$ = 0.2289<br>**** $F$ (1, 7) = 95.55, $p$ < 0.0001 | |
| **6G** | 2-Way RM ANOVA | day x light phase<br>day<br>light phase | $F$ (1.924, 10.10) = 1.518, $p$ = 0.2646<br>* $F$ (1.88, 13.16) = 4.877, $p$ = 0.0276<br>$F$ (1. 7) = 1.826, $p$ = 0.2186 | *<u>Baseline vs Kynurenine:</u>*<br>* day 2 (light phase): $p$ < 0.05<br>* day 4 (light phase): $p$ < 0.05<br>* day 6 (light phase): $p$ < 0.05<br>* washout (dark phase): $p$ < 0.05 |

| Supplemental Figure | Statistical Test | Interaction<br>Main Effects | Results | Fisher's LSD Posthoc Results: |
|---|---|---|---|---|
| S4B | 2-way ANOVA | pregnancy x kynurenine<br>pregnancy<br>kynurenine | $F$ (1, 28) = 1.215, $p$ = 0.2797<br>**** $F$ (1, 28) = 37.14, $p$ < 0.0001<br>$F$ (1, 28) = 2.009, $p$ = 0.1674 | ** Control vs ECon: $p$ < 0.01<br>**** Kynurenine vs EKyn: $p$ < 0.0001 |
| S4C | 2-way ANOVA | pregnancy x kynurenine<br>pregnancy<br>kynurenine | $F$ (1, 27) = 3.007, $p$ = 0.0943<br>* $F$ (1, 27) = 4.919, $p$ = 0.0352<br>**** $F$ (1, 27) = 121.6, $p$ < 0.0001 | * Kynurenine vs EKyn: $p$ < 0.05 |
| S4D | 2-way ANOVA | pregnancy x kynurenine<br>pregnancy<br>kynurenine | $F$ (1, 27) = 0.8115, $p$ = 0.3756<br>** $F$ (1, 27) = 12.21, $p$ = 0.0017<br>$F$ (1, 27) = 30.95, $p$ < 0.0001 | ** Kynurenine vs EKyn: $p$ < 0.01 |
| S4E | 2-way ANOVA | pregnancy x kynurenine<br>pregnancy<br>kynurenine | $F$ (1, 18) = 0.0118, $p$ = 0.9147<br>* $F$ (1, 18) = 6.266, $p$ = 0.0222<br>* $F$ (1, 18) = 6.181, $p$ = 0.0230 | |
| S5B | 2-way ANOVA | pregnancy x kynurenine<br>pregnancy<br>kynurenine | $F$ (1, 24) = 0.003868, $p$ = 0.9509<br>$F$ (1, 24) = 0.9086, $p$ = 0.35<br>**** $F$ (1, 24) = 33.7, $p$ < 0.0001 | |
| S5C | 2-way ANOVA | pregnancy x kynurenine<br>pregnancy<br>kynurenine | * $F$ (1, 27) = 7.351, $p$ = 0.0115<br>**** F (1, 27) = 25.74, $p$ < 0.0001<br>* F (1, 27) = 6.583, $p$ = 0.0162 | * Kynurenine vs EKyn: $p$ < 0.001 |
| S5D | 2-way ANOVA | pregnancy x kynurenine<br>pregnancy<br>kynurenine | $F$ (1, 30) = 3.10, $p$ = 0.0885<br>* $F$ (1, 30) = 4.817, $p$ = 0.0361<br>**** $F$ (1, 30) = 64.25, $p$ < 0.0001 | * Kynurenine vs EKyn: $p$ < 0.05 |
| S5E | 2-way ANOVA | pregnancy x kynurenine<br>pregnancy<br>kynurenine | $F$ (1, 25) = 0.6055, $p$ = 0.4438<br>$F$ (1, 25) = 0.5215, $p$ = 0.4769<br>** $F$ (1, 25) = 10.20, $p$ = 0.0038 | |
| S5F | 2-way ANOVA | pregnancy x kynurenine<br>pregnancy<br>kynurenine | $F$ (1, 22) = 1.290, $p$ = 0.2682<br>$F$ (1, 22) = 0.04489, $p$ = 0.8342<br>** $F$ (1, 22) = 8.293, $p$ = 0.0087 | |
| S5G | 2-way ANOVA | pregnancy x kynurenine<br>pregnancy<br>kynurenine | $F$ (1, 24) = 0.006697, $p$ = 0.9355<br>** $F$ (1, 24) = 10.09, $p$ = 0.0041<br>* $F$ (1, 24) = 4.668, $p$ = 0.0409 | * Control vs ECon: $p$ < 0.05<br>* Kynurenine vs EKyn: $p$ < 0.05 |
| S5H | 2-way ANOVA | pregnancy x kynurenine<br>pregnancy<br>kynurenine | $F$ (1, 23) = 2.340, $p$ = 0.1397<br>$F$ (1, 23) = 0.2446, $p$ = 0.6256<br>* $F$ (1, 23) = 7.278, $p$ = 0.0128 | |
| S5I | 2-way ANOVA | pregnancy x kynurenine<br>pregnancy<br>kynurenine | * $F$ (1, 27) = 6.290, $p$ = 00185<br>**** $F$ (1, 27) = 24.47, $p$ < 0.0001<br>** $F$ (1, 27) = 7.973, $p$ = 0.0088 | *** Kynurenine vs Ekyn: $p$ < 0.001 |
| S5J | 2-way ANOVA | pregnancy x kynurenine<br>pregnancy<br>kynurenine | $F$ (1, 28) = 3.124, $p$ = 0.0881<br>**** $F$ (1, 28) = 24.63, $p$ < 0.0001<br>** $F$ (1, 28) = 11.29, $p$ = 0.0023 | * Control vs ECon: $p$ < 0.05<br>**** Kynurenine vs EKyn: $p$ < 0.0001 |
| S5K | 2-way ANOVA | pregnancy x kynurenine<br>pregnancy<br>kynurenine | $F$ (1, 21) = 3.348, $p$ = 0.0815<br>* $F$ (1, 21) = 4.772, $p$ = 0.0404<br>** $F$ (1, 21) = 11.43, $p$ = 0.0028 | ** Kynurenine vs EKyn: $p$ < 0.01 |
| S6A | 2-way RM ANOVA | cohort x light phase<br>cohort<br>light phase | $F$ (2, 20) = 0.2436, $p$ = 0.7861<br>$F$ (2, 20) = 0.2042, $p$ = 0.8169<br>**** $F$ (1, 20) = 947, $p$ < 0.0001 | |
| S6B | 2-way RM ANOVA | cohort x light phase<br>cohort<br>light phase | $F$ (2, 20) = 0.4193, $p$ = 0.6632<br>$F$ (2, 20) = 0.7152, $p$ = 0.5012<br>**** $F$ (1, 20) = 194, $p$ < 0.0001 | |
| S6C | 2-way RM ANOVA | cohort x light phase<br>cohort<br>light phase | $F$ (2, 20) = 0.8653, $p$ = 0.4361<br>$F$ (2, 20) = 0.8052, $p$ = 0.4610<br>**** $F$ (1, 20) = 112.3, $p$ < 0.0001 | |
| S6D | 2-way RM ANOVA | cohort x light phase<br>cohort<br>light phase | $F$ (2, 20) = 0.1261, $p$ = 0.8822<br>$F$ (2, 20) = 0.7816, $p$ = 0.4712<br>**** $F$ (1, 20) = 1034, $p$ < 0.0001 | |
| S6E | 2-way RM ANOVA | cohort x light phase<br>cohort<br>light phase | $F$ (2, 20) = 0.049, $p$ = 0.9522<br>$F$ (2, 20) = 1.887, $p$ = 0.1775<br>**** $F$ (1, 20) = 306.2, $p$ < 0.0001 | |
| S6F | 2-way RM ANOVA | cohort x light phase<br>cohort<br>light phase | $F$ (2, 20) = 0.8297, $p$ = 0.4507<br>$F$ (2, 20) = 0.8958, $p$ = 0.424<br>**** $F$ (1, 20) = 107.3, $p$ < 0.0001 | |

| Supplemental Figure | Statistical Test | Interaction<br>Main Effects | Results | Fisher's LSD Posthoc Results: |
|---|---|---|---|---|
| S6G | 2-way RM ANOVA | cohort x light phase<br>cohort<br>light phase | *F* (2, 20) = 0.5091, *p* = 0.6086<br>*F* (2, 20) = 0.9843, *p* = 0.3911<br>**** *F* (1, 20) = 410.8, *p* < 0.0001 | |
| S6H | 2-way RM ANOVA | cohort x light phase<br>cohort<br>light phase | *F* (2, 20) = 0.4466, *p* = 0.6460<br>*F* (2, 20) = 0.5775, *p* = 0.5704<br>**** *F* (1, 20) = 458.8, *p* < 0.0001 | |
| S6I | 2-way RM ANOVA | cohort x light phase<br>cohort<br>light phase | *F* (2, 20) = 0.0886, *p* = 0.9156<br>*F* (2, 20) = 1.4, *p* = 0.2698<br>*** *F* (1, 20) = 15.25, *p* = 0.0009 | |
| S7A | 2-way RM ANOVA | pregnancy x light phase<br>pregnancy<br>light phase | **** *F* (2.988, 17.18) = 35.12, *p* < 0.0001<br>**** *F* (1.865, 13.05) = 24.68, *p* < 0.0001<br>**** *F* (1, 7) = 619.4, *p* < 0.0001 | *Baseline vs ECon:*<br>* PD 5 (light phase): *p* < 0.01<br>*** ED 16 (dark phase): *p* < 0.001<br>**** ED 18 (dark phase): *p* < 0.0001<br>**** ED 20 (dark phase): *p* < 0.0001<br>** PD 5 (dark phase): *p* < 0.01 |
| S7B | 2-way RM ANOVA | pregnancy x light phase<br>pregnancy<br>light phase | *F* (2.449, 14.08) = 2.361, *p* = 0.1232<br>* *F* (2.520, 17.64) = 4.110, *p* = 0.0271<br>**** *F* (1,7) = 163.3, *p* < 0.0001 | *Baseline vs ECon:*<br>* ED 20 (dark phase): *p* < 0.05<br>* PD 5 (dark pahse): *p* < 0.05 |
| S7C | 2-way RM ANOVA | pregnancy x light phase<br>pregnancy<br>light phase | * *F* (2.709, 15.58) = 4.413, *p* = 0.0221<br>*F* (2.759, 19.31) = 2.999, *p* = 0.0595<br>*** *F* (1, 7) = 29.59, *p* = 0.001 | *Baseline vs ECon:*<br>* ED 20 (dark phase): *p* < 0.05<br>* PD 5 (dark pahse): *p* < 0.05 |
| S7D | 2-way RM ANOVA | pregnancy x light phase<br>pregnancy<br>light phase | *F* (1,706, 7.252) = 3.221, *p* = 0.1037<br>* *F* (2.190, 15.33) = 3.997, *p* = 0.0371<br>**** *F* (1, 7) = 145.9, *p* < 0.0001 | *Baseline vs ECon:*<br>* ED 16 (dark phase): *p* < 0.05<br>* ED 20 (dark pahse): *p* < 0.05 |
| S7E | 2-way RM ANOVA | pregnancy x light phase<br>pregnancy<br>light phase | * *F* (2.074, 11.40) = 6.236, *p* = 0.0142<br>**** *F* (2.344, 16.41) = 66.95, *p* < 0.0001<br>*** *F* (1, 7) = 132.5, *p* < 0.0001 | *Baseline vs ECon:*<br>* ED 18 (light phase): *p* < 0.05<br>**** PD 5 (light pahse): *p* < 0.0001<br>**** ED 16 (dark phase): *p* < 0.0001<br>*** ED 18 (dark phase): *p* < 0.001<br>** ED 20 (dark phase): *p* < 0.01<br>** PD5 (dark phase): *p* < 0.01 |
| ECon Wake to NREM | 2-way RM ANOVA | day x light phase<br>day<br>light phase | *F* (2.359, 10.62) = 2.952, *p* = 0.0899<br>* *F* (2.348, 14.09) = 4.285, *p* = 0.0305<br>**** *F* (2.359, 10.62) = 2.952, *p* < 0.0001 | *Baseline vs ECon:*<br>* ED 20 (dark phase): *p* < 0.05<br>** PD 5 (dark phase): *p* < 0.01 |
| ECon NREM to Wake | 2-way RM ANOVA | day x light phase<br>day<br>light phase | *F* (2.070, 9.313) = 1.975, *p* = 0.1924<br>** *F* (2.828, 16.97) = 6.125, *p* = 0.0057<br>****F* (1, 6) = 47.77, *p* = 0.0005 | *Baseline vs ECon:*<br>* ED 16 (light phase): *p* < 0.05<br>* ED 18 (light phase): *p* < 0.05<br>* ED 18 (dark phase): *p* < 0.05 |
| ECon NREM to REM | 2-way RM ANOVA | day x light phase<br>day<br>light phase | ** *F* (1.707, 7.680) = 11.59, *p* = 0.0057<br>*F* (1.744, 10.46) = 1.707, *p* = 0.2279<br>**** *F* (1, 6) = 266.6, *p* < 0.0001 | *Baseline vs ECon:*<br>* PD 5 (light phase): *p* < 0.05<br>* ED 16 (dark phase): *p* < 0.05<br>** ED 18 (dark phase): *p* < 0.01<br>* ED 20 (dark phase): *p* < 0.05<br>* PD 5 (dark phase): *p* < 0.05 |
| ECon REM to Wake | 2-way RM ANOVA | day x light phase<br>day<br>light phase | ** *F* (1.615, 7.269) = 11.66, *p* = 0.0067<br>*F* (1.568, 9.408) = 1.701, *p* = 0.2322<br>**** *F* (1, 6) = 243.9, *p* < 0.0001 | *Baseline vs ECon:*<br>** PD 5 (light phase): *p* < 0.01<br>* ED 16 (dark phase): *p* < 0.05<br>** ED 18 (dark phase): *p* < 0.01<br>* ED 20 (dark phase): *p* < 0.05<br>* PD 5 (dark phase): *p* < 0.05 |
| ECon REM to NREM | 2-way RM ANOVA | day x light phase<br>day<br>light phase | *F* (2.540, 34.29) = 1.112, *p* = 0.3512<br>*F* (2.047, 27.63) = 2.219, *p* = 0.1267<br>**** *F* (1, 54) = 29.81, *p* < 0.0001 | |
| S8A | 2-way RM ANOVA | pregnancy x light phase<br>pregnancy<br>light phase | *** *F* (1.972, 10.85) = 21.86, *p* = 0.0002<br>*** *F* (2.941, 17.64) = 9.815, *p* = 0.0005<br>**** *F* (1, 6) = 377.1, *p* < 0.0001 | *Baseline vs EKyn:*<br>ED 20 (light phase): *p* < 0.01<br>*** ED 16 (dark phase): *p* < 0.001<br>**** ED 18 (dark phase): *p* < 0.0001<br>** ED 20 (dark phase): *p* < 0.01<br>**** PD 5 (dark phase): *p* < 0.0001 |

| Supplemental Figure | Statistical Test | Interaction<br>Main Effects | Results | Fisher's LSD Posthoc Results: |
|---|---|---|---|---|
| S8B | 2-way RM ANOVA | pregnancy x light phase<br>pregnancy<br>light phase | * $F$ (1.783, 9.808) = 4.797, $p$ = 0.0382<br>** $F$ (2.612, 15.67) = 9.239, $p$ = 0.0013<br>**** $F$ (1, 6) = 133.6, $p$ < 0.0001 | _Baseline vs EKyn:_<br>* ED 16 (dark phase): $p$ < 0.05<br>** PD 5 (dark phase): $p$ < 0.01 |
| S8C | 2-way RM ANOVA | pregnancy x light phase<br>pregnancy<br>light phase | ** $F$ (2.108, 11.60) = 7.136, $p$ = 0.0089<br>$F$ (2.093, 12.56) = 3.555, $p$ = 0.0582<br>**** $F$ (1, 6) = 94.27, $p$ < 0.0001 | _Baseline vs EKyn:_<br>** ED 16 (dark phase): $p$ < 0.01<br>** ED 18 (dark phase): $p$ < 0.01<br>** PD 5 (dark phase): $p$ < 0.01 |
| S8D | 2-way RM ANOVA | pregnancy x light phase<br>pregnancy<br>light phase | * $F$ (2.729, 17.06) = 4.347, $p$ = 0.0213<br>$F$ (2.297, 16.08) = 2.650, $p$ = 0.0955<br>**** $F$ (1, 7) = 96.22, $p$ < 0.0001 | |
| S8E | 2-way RM ANOVA | pregnancy x light phase<br>pregnancy<br>light phase | **** $F$ (2.457, 15.97) = 19.34, $p$ < 0.0001<br>**** $F$ (1.518, 10.62) = 66.93, $p$ < 0.0001<br>**** $F$ (1, 7) = 205.4, $p$ < 0.0001 | _Baseline vs EKyn:_<br>* ED 20 (light phase): $p$ < 0.05<br>**** PD 5 (light pahse): $p$ < 0.0001<br>**** ED 16 (dark phase): $p$ < 0.0001<br>**** ED 18 (dark phase): $p$ < 0.0001<br>*** ED 20 (dark phase): $p$ < 0.001<br>** PD5 (dark phase): $p$ < 0.01 |
| EKyn Wake to NREM | 2-way RM ANOVA | day x light phase<br>day<br>light phase | $F$ (1.914, 11) = 3.831, $p$ = 0.0560<br>*** $F$ (3.032, 18.19) = 8.526, $p$ = 0.0009<br>**** $F$ (1. 6) = 111.5, $p$ < 0.0001 | _Baseline vs EKyn:_<br>* ED 16 (light phase): $p$ < 0.05<br>** PD 5 (dark phase): $p$ < 0.01 |
| EKyn NREM to Wake | 2-way RM ANOVA | day x light phase<br>day<br>light phase | $F$ (2.009, 11.55) = 0.6775, $p$ = 0.5276<br>**** $F$ (2.819, 16.92) = 15.90, $p$ < 0.0001<br>** $F$ (1, 6) = 25.76, $p$ = 0.0023 | _Baseline vs EKyn:_<br>* ED 16 (light phase): $p$ < 0.05<br>** PD 5 (light phase): $p$ < 0.01<br>* PD 5 (dark phase): $p$ < 0.05 |
| EKyn NREM to REM | 2-way RM ANOVA | day x light phase<br>day<br>light phase | **** $F$ (2.454, 14.11) = 17.51, $p$ < 0.0001<br>$F$ (1.635, 9.813) = 3.435, $p$ = 0.0804<br>**** $F$ (1, 6) = 118.6, $p$ < 0.0001 | _Baseline vs EKyn:_<br>** PD 5 (light phase): $p$ < 0.01<br>* ED 16 (dark phase): $p$ < 0.05<br>** ED 18 (dark phase): $p$ < 0.01<br>* ED 20 (dark phase): $p$ < 0.05<br>**** PD 5 (dark phase): $p$ < 0.0001 |
| EKyn REM to Wake | 2-way RM ANOVA | day x light phase<br>day<br>light phase | **** $F$ (2.407, 13.84) = 18.50, $p$ < 0.0001<br>* $F$ (1.794, 10.77) = 5.989, $p$ = 0.0199<br>**** $F$ (1, 6) = 117.6, $p$ < 0.0001 | _Baseline vs EKyn:_<br>*** PD 5 (light phase): $p$ < 0.001<br>* ED 16 (dark phase): $p$ < 0.05<br>** ED 18 (dark phase): $p$ < 0.01<br>* ED 20 (dark phase): $p$ < 0.05<br>**** PD 5 (dark phase): $p$ < 0.0001 |
| EKyn REM to NREM | 2-way RM ANOVA | day x light phase<br>day<br>light phase | $F$ (1.387, 7.976) = 0.5387, $p$ = 0.5410<br>*** $F$ (1.949, 11.70) = 14.87, $p$ = 0.0006<br>$F$ (1. 6) = 5.778, $p$ = 0.0530 | _Baseline vs EKyn:_<br>* ED 20 (light phase): $p$ < 0.05<br>** PD 5 (light phase): $p$ < 0.01<br>* ED 20 (dark phase): $p$ < 0.05<br>* PD 5 (dark phase): $p$ < 0.05 |
| S9A | 2-way ANOVA | EKyn x light phase<br>EKyn<br>light phase | $F$ (1, 13) = 1.846, $p$ = 0.1973<br>$F$ (1, 13) = 3.280, $p$ = 0.0933<br>**** $F$ (1, 13) = 431.8, $p$ < 0.0001 | |
| S9B | 2-way ANOVA | EKyn x light phase<br>EKyn<br>light phase | $F$ (1, 13) = 0.2713, $p$ = 0.6112<br>$F$ (1, 13) = 0.6418, $p$ = 0.4375<br>**** $F$ (1, 13) = 308.3, $p$ < 0.0001 | |
| S9C | 2-way ANOVA | EKyn x light phase<br>EKyn<br>light phase | $F$ (1, 13) = 2.510, $p$ = 0.1371<br>$F$ (1, 13) = 2.402, $p$ = 0.1452<br>**** $F$ (1, 13) = 102,7, $p$ < 0.0001 | |
| S9D | 2-way ANOVA | EKyn x light phase<br>EKyn<br>light phase | $F$ (1, 13) = 3.736, $p$ = 0.0753<br>$F$ (1, 13) = 0.03289, $p$ = 0.8589<br>**** $F$ (1, 13) = 488.6, $p$ < 0.0001 | |
| S9E | 2-way ANOVA | EKyn x light phase<br>EKyn<br>light phase | $F$ (1, 13) = 0.044, $p$ = 0.8362<br>$F$ (1, 13) = 1.370, $p$ = 0.2629<br>**** $F$ (1, 13) = 356.8, $p$ < 0.0001 | |
| S9F | 2-way ANOVA | EKyn x light phase<br>EKyn<br>light phase | $F$ (1, 13) = 2.349, $p$ = 0.1493<br>$F$ (1, 13) = 0.0102, $p$ = 0.9211<br>** $F$ (1, 13) = 10.46, $p$ = 0.0065 | |
| S9G | 2-way ANOVA | EKyn x light phase<br>EKyn<br>light phase | $F$ (1, 13) = 0.1288, $p$ = 0.7254<br>$F$ (1, 13) = 0.1797, $p$ = 0.6786<br>**** $F$ (1, 13) = 82.44, $p$ < 0.0001 | |

| Supplemental Figure | Statistical Test | Interaction<br>Main Effects | Results | Fisher's LSD Posthoc Results: |
|---|---|---|---|---|
| S9H | 2-way ANOVA | EKyn x light phase<br>EKyn<br>light phase | $F$ (1, 13) = 2.963, $p$ = 0.1089<br>$F$ (1, 13) = 1.091, $p$ = 0.3153<br>**** $F$ (1, 13) = 214.8, $p$ < 0.0001 | |
| S9I | 2-way ANOVA | EKyn x light phase<br>EKyn<br>light phase | $F$ (1, 13) = 0.03264, $p$ = 0.8594<br>$F$ (1, 13) = 4.025, $p$ = 0.0661<br>$F$ (1, 13) = 1.298, $p$ = 0.2752 | |
| S10A | 2-way ANOVA | EKyn x light phase<br>EKyn<br>light phase | $F$ (1, 12) = 0.8888, $p$ = 0.3644<br>$F$ (1, 12) = 1.883, $p$ = 0.1951<br>**** $F$ (1, 12) = 119.7, $p$ < 0.0001 | |
| S10B | 2-way ANOVA | EKyn x light phase<br>EKyn<br>light phase | $F$ (1, 12) = 0.3512, $p$ = 0.5644<br>$F$ (1, 12) = 0.0197, $p$ = 0.891<br>** $F$ (1, 12) = 12.07, $p$ = 0.0046 | |
| S10C | 2-way ANOVA | EKyn x light phase<br>EKyn<br>light phase | $F$ (1, 12) = 0.4593, $p$ = 0.5108<br>$F$ (1, 12) = 0.5642, $p$ = 0.4671<br>$F$ (1, 12) = 3.173, $p$ = 0.1002 | |
| S10D | 2-way ANOVA | EKyn x light phase<br>EKyn<br>light phase | $F$ (1, 12) = 0.5645, $p$ = 0.4669<br>$F$ (1, 12) = 1.184, $p$ = 0.2977<br>**** $F$ (1, 12) = 171.4, $p$ < 0.0001 | |
| S10E | 2-way ANOVA | EKyn x light phase<br>EKyn<br>light phase | $F$ (1, 12) = 0.2723, $p$ = 0.6133<br>$F$ (1, 12) = 0.06851, $p$ = 0.7980<br>** $F$ (1, 12) = 14.52, $p$ = 0.0025 | |
| S10F | 2-way ANOVA | EKyn x light phase<br>EKyn<br>light phase | $F$ (1, 12) = 0.7192, $p$ = 0.4130<br>$F$ (1, 12) = 0.008675, $p$ = 0.9273<br>* $F$ (1, 12) = 6.749, $p$ = 0.0233 | |
| S10G | 2-way ANOVA | EKyn x light phase<br>EKyn<br>light phase | $F$ (1, 12) = 0.04339, $p$ = 0.8385<br>$F$ (1, 12) = 0.7520, $p$ = 0.1045<br>** $F$ (1, 12) = 10.71, $p$ = 0.0067 | |
| S10H | 2-way ANOVA | EKyn x light phase<br>EKyn<br>light phase | $F$ (1, 12) = 0.000289, $p$ = 0.9867<br>$F$ (1, 12) = 1.476, $p$ = 0.2478<br>* $F$ (1, 12) = 4.812, $p$ = 0.0487 | |
| S10I | 2-way ANOVA | EKyn x light phase<br>EKyn<br>light phase | $F$ (1, 12) = 0.1505, $p$ = 0.7049<br>$F$ (1, 12) = 2.866, $p$ = 0.1163<br>$F$ (1, 12) = 3.731, $p$ = 0.774 | |
| S11A | 2-way RM ANOVA | day x light phase<br>day<br>light phase | * $F$ (2.208, 12.70) = 5.432, $p$ = 0.0177<br>$F$ (2.158, 15.10) = 0.7129, $p$ = 0.5161<br>**** $F$ (1, 7) = 330.3, $p$ < 0.0001 | *Baseline vs Kynurenine:*<br>day 2 (light phaes): $p$ < 0.01<br>day 4 (light phase): $p$ < 0.01<br>day 6 (light phase): $p$ < 0.05 |
| S11B | 2-way RM ANOVA | day x light phase<br>day<br>light phase | $F$ (1.973, 11.84) = 0.3842, $p$ = 0.6865<br>$F$ (2.547, 17.20) = 1.499, $p$ = 0.2513<br>**** $F$ (1, 7) = 85.9, $p$ < 0.0001 | |
| S11C | 2-way RM ANOVA | day x light phase<br>day<br>light phase | $F$ (2.058, 12.35) = 0.5918, $p$ = 0.5728<br>$F$ (2.319, 16.23) = 1.026, $p$ = 0.3906<br>*** $F$ (1, 7) = 46.17, $p$ = 0.003 | |
| S11D | 2-way RM ANOVA | day x light phase<br>day<br>light phase | $F$ (1.987, 12.42) = 2.431, $p$ = 0.1287<br>$F$ (2.056, 14.39) = 2.141, $p$ = 0.1527<br>** $F$ (1, 7) = 23.43, $p$ = 0.0019 | |
| S11E | 2-way RM ANOVA | day x light phase<br>day<br>light phase | $F$ (2.408, 10.84) = 1.225, $p$ = 0.3392<br>$F$ (2.295, 16.06) = 1.335, $p$ = 0.2937<br>**** $F$ (1, 7) = 115.8, $p$ < 0.0001 | |
| S12A | 2-way RM ANOVA | light phase:<br>kynurenine x frequency<br>kynurenine<br>frequency | $F$ (1.331, 7.793) = 2.5774, $p$ = 0.1459<br>$F$ (1. 7) = 0.4560, $p$ = 0.5212<br>**** $F$ (1.191, 8.336) = 43.77, $p$ < 0.0001 | |
| | | dark phase:<br>kynurenine x frequency<br>kynurenine<br>frequency | $F$ (1.431, 6.746) = 2.118, $p$ = 0.1938<br>$F$ (1, 7) = 0.1356, $p$ = 0.7235<br>** $F$ (1.348, 9.433) = 18.56, $p$ = 0.0011 | |

| Supplemental Figure | Statistical Test | Interaction<br>Main Effects | Results | Fisher's LSD Posthoc Results: |
|---|---|---|---|---|
| S12B | 2-way RM ANOVA | light phase:<br>kynurenine x frequency<br>kynurenine<br>frequency | * $F$ (1.576, 9.260) = 4.868, $p$ = 0.0419<br>$F$ (1, 7) = 0.5667, $p$ = 0.4761<br>**** $F$ (1.746, 12.22) = 51.79, $p$ < 0.0001 | *Kynurenine vs Baseline:*<br>* 7.5 Hz: $p$ < 0.05 |
| | | dark phase:<br>kynurenine x frequency<br>kynurenine<br>frequency | $F$ (1.303, 6.191) = 2.186, $p$ = 0.1911<br>$F$ (1, 7) = 0.2537, $p$ = 0.63<br>**** $F$ (1.999, 13.99) = 35.09, $p$ < 0.0001 | |
| S12C | 2-way RM ANOVA | light phase:<br>kynurenine x frequency<br>kynurenine<br>frequency | $F$ (0.001626, 0.007664) = 0.7571, $p$ = 0.1762<br>$F$ (1, 7) = 0.1044, $p$ = 0.756<br>*** $F$ (1.048, 7.338) = 35.58, $p$ = 0.0004 | |
| | | dark phase:<br>kynurenine x frequency<br>kynurenine<br>frequency | $F$ (0.09135, 0.4306) = 3.487, $p$ = 0.1808<br>$F$ (1, 7) = 0.523, $p$ = 0.4816<br>** $F$ (1.054, 7.375) = 16.61, $p$ = 0.004 | |
| S12D | 2-way RM ANOVA | light phase:<br>kynurenine x frequency<br>kynurenine<br>frequency | $F$ (1.692, 8.035) = 2.699, $p$ = 0.1308<br>$F$ (1, 7) = 0.6122, $p$ = 0.4596<br>**** $F$ (1.457, 10.20) = 36.11, $p$ < 0.0001 | |
| | | dark phase:<br>kynurenine x frequency<br>kynurenine<br>frequency | $F$ (1.474, 6.999) = 2.199, $p$ = 0.1834<br>$F$ (1. 7) = 0.0883, $p$ = 0.775<br>*** $F$ (1.507, 10.55) = 28.44, $p$ = 0.0001 | |
| Kynurenine Wake to NREM | 2-way RM ANOVA | day x light phase<br>day<br>light phase | $F$ (1.618, 10.51) = 1.624, $p$ = 0.2408<br>$F$ (2.379, 16.65) = 1.579, $p$ = 0.2348<br>**** $F$ (1, 7) = 178, $p$ < 0.0001 | |
| Kynurenine NREM to Wake | 2-way RM ANOVA | day x light phase<br>day<br>light phase | $F$ (2.148, 13.96) = 1.463, $p$ = 0.2659<br>$F$ (2.464, 17.25) = 0.5303, $p$ = 0.6338<br>** $F$ (1, 7) = 17.48, $p$ = 0.0041 | |
| Kynurenine NREM to REM | 2-way RM ANOVA | day x light phase<br>day<br>light phase | $F$ (1.650, 9.9) = 2.069, $p$ = 0.1802<br>$F$ (2.286, 16) = 1.935, $p$ = 0.1733<br>**** $F$ (1. 7) = 85.96, $p$ = 0.1733 | |
| Kynurenine REM to Wake | 2-way RM ANOVA | day x light phase<br>day<br>light phase | $F$ (1.484, 9.647) = 2.074, $p$ = 0.1817<br>$F$ (2.748, 19.24) = 1.637, $p$ = 0.2158<br>**** $F$ (1, 7) = 87.84, $p$ < 0.0001 | |
| Kynurenine REM to NREM | 2-way RM ANOVA | day x light phase<br>day<br>light phase | $F$ (2.963, 19.26) = 1.484, $p$ = 0.2505<br>$F$ (2.256, 15.79) = 2.293, $p$ = 0.1291<br>* $F$ (1. 7) = 10.65, $p$ = 0.0138 | |
| Kynurenine NREM Onset | 1-way RM ANOVA | day | $F$ (1.254, 8.148) = 2.645, $p$ = 0.1394 | |
| ECon NREM Onset | 1-way RM ANOVA | day | $F$ (1.928, 13.98) = 2.397, $p$ = 0.1286 | |
| EKyn NREM Onset | 1-way RM ANOVA | day | $F$ (2.542, 15.25) = 1.329, $p$ = 0.2990 | |
| Kynurenine REM Onset | 1-way RM ANOVA | day | $F$ (1.578, 14.21) = 2.187, $p$ = 0.1548 | |
| ECon REM Onset | 1-way RM ANOVA | day | $F$ (1.875, 10.31) = 1.145, $p$ = 0.3517 | |
| EKyn REM Onset | 1-way RM ANOVA | day | $F$ (1.278, 7.667) = 2.138, $p$ = 0.1851 | |
